# Periodic propagating waves coordinate RhoGTPase network dynamics at the leading and trailing edges during cell migration

**DOI:** 10.1101/2020.03.09.984054

**Authors:** Alfonso Bolado-Carrancio, Oleksii S. Rukhlenko, Elena Nikonova, Mikhail A. Tsyganov, Anne Wheeler, Amaya Garcia Munoz, Walter Kolch, Alex von Kriegsheim, Boris N. Kholodenko

**Affiliations:** Edinburgh Cancer Research Centre, Institute of Genetics and Molecular Medicine, University of Edinburgh, Edinburgh, EH4 2XR, United Kingdom; Systems Biology Ireland, School of Medicine and Medical Science, University College Dublin, Belfield, Dublin 4, Ireland; Institute of Theoretical and Experimental Biophysics, Pushchino, Moscow Region, Russia; Conway Institute of Biomolecular & Biomedical Research, University College Dublin, Belfield, Dublin 4, Ireland; Department of Pharmacology, Yale University School of Medicine, New Haven, USA

## Abstract

Migrating cells need to coordinate distinct leading and trailing edge dynamics but the underlying mechanisms are unclear. Here, we combine experiments and mathematical modeling to elaborate the minimal autonomous biochemical machinery necessary and sufficient for this dynamic coordination and cell movement. RhoA activates Rac1 via DIA and inhibits Rac1 via ROCK, while Rac1 inhibits RhoA through PAK. Our data suggest that in motile, polarized cells, RhoA–ROCK interactions prevail at the rear whereas RhoA-DIA interactions dominate at the front where Rac1/Rho oscillations drive protrusions and retractions. At the rear, high RhoA and low Rac1 activities are maintained until a wave of oscillatory GTPase activities from the cell front reaches the rear, inducing transient GTPase oscillations and RhoA activity spikes. After the rear retracts, the initial GTPase pattern resumes. Our findings show how periodic, propagating GTPase waves coordinate distinct GTPase patterns at the leading and trailing edge dynamics in moving cells.

## Introduction

Cell migration relies on the coordination of actin dynamics at the leading and the trailing edges ^1^. During the mesenchymal type of cell migration, protrusive filamentous actin (F-actin) is cyclically polymerised/depolymerised at the cell’s leading edge, whereas the contractile, actomyosin-enriched trailing edge forms the rear. The leading edge protrudes and retracts multiple times, until the protrusions, known as lamellipodia, are stabilized by adhering to the extracellular matrix ^2^. Subsequently, the cell rear detaches and contracts allowing the cell body to be pulled towards the front. Core biochemical mechanisms of this dynamic cycle are governed by the Rho family of small GTPases ^3^. Two members of this family, Ras homolog family member A (RhoA) and Ras-related C3 botulinum toxin substrate 1 (Rac1), control protrusions and retractions at the leading edge as well as the contractility at the rear ^4–6^. Rho GTPases cycle between an active, GTP-loaded ‘on’ state and an inactive, GDP-loaded ‘off’ state. Switches between on and off states are tightly regulated by (i) guanine nucleotide exchange factors (GEFs) that facilitate GDP/GTP exchange thereby activating GTPases and (ii) GTPase activating proteins (GAPs) that stimulate GTP hydrolysis and transition to a GDP-bound state.

A canonic description of mesenchymal cell migration portrays mutually separated zones of Rac1-GTP and RhoA-GTP in polarized cells where Rac1-GTP dominates at the leading edge and RhoA-GTP dominates at the contracted cell rear ^7–13^. This distinct distribution of RhoA and Rac1 activities along polarized cells is explained by a mutual antagonism of RhoA and Rac1 ^14,15^ mediated by their downstream effectors ^16–18^. The Rac1 effector, p21 associated kinase (PAK), phosphorylates and inhibits multiple RhoA-specific GEFs, including p115-RhoGEF, GEF-H1 and Net1 ^16,19,20^. In addition, active Rac1 binds and activates p190RhoGAP, which decreases RhoA activity ^16^. In turn, RhoA-GTP recruits the Rho-associated kinase (ROCK), which phosphorylates and activates Rac-specific GAPs, such as FilGAP and ArhGAP22, thereby inhibiting Rac1 ^16,21,22^. This mutual inhibition of RhoA and Rac1 may lead to a bistable behavior where a system can switch between two stable steady states, in which GTPase activities alternate between high and low values ^14,23^. The existence of bistable switches is supported by experiments, where inhibition of the Rac1 effector PAK maintains both high RhoA and low Rac1 activities and associated morphological changes even after the inhibition is released ^17^.

At the same time, RhoA and Rac1 do not behave antagonistically at the leading edge of migrating cells. Here, RhoA activation is rapidly followed by Rac1 activation, tracking a protrusion-retraction cycle ^6^. This Rac1 activation at the leading edge is mediated by the downstream RhoA effector, Diaphanous related formin-1 (DIA), that was shown to localize to the membrane ruffles of motile cells ^24,25^. Thus, in contrast to the RhoA effector ROCK, which inhibits Rac1 in the other cell segments, the RhoA effector DIA can stimulate Rac1 activity at the leading edge.

If at the leading edge RhoA activates Rac1 but Rac1 inhibits RhoA, this intertwined network circuitry of positive and negative loops will force the network to periodically change RhoA and Rac1 activities, giving rise to self-perpetuating oscillations with a constant amplitude and frequency ^23,26^. By contrast, at the trailing edge and cell body, the mutual RhoA and Rac1 inhibition results in the maintenance of a (quasi)steady state with high RhoA activity and low Rac1 activity. But, how can these different dynamics coexist? More importantly, how are these dynamics coordinated within the cell?

Here, we first elucidated the spatial profiles of RhoA-Rac1 interactions in motile MDA-MB-231 breast cancer cells. Using proximity ligation assays (PLA), we show that the concentration of complexes formed by RhoA and its downstream effectors DIA and ROCK depends on the spatial location along the longitudinal axis of polarized cells. RhoA primarily interacts with DIA at the cell leading edge, whereas RhoA - ROCK interactions are the strongest at the cell rear. Based on these findings, we built a mathematical model to analyze RhoA-Rac1 signaling in space and time. The model predicts and the experiments corroborate that at the cell front the GTPase network exhibits oscillatory behavior with high average Rac1-GTP, whereas at the cell rear there is a (quasi)steady state with high RhoA-GTP and low Rac. The front and rear are connected by periodic, propagating GTPase waves. When the wave reaches the rear, RhoA-GTP transiently oscillates and then, following the rear retraction, the GTPase network dynamic pattern returns to the original state. Our model and experimental results show how different GTPase dynamics at the leading edge and the rear can govern distinct cytoskeleton processes and how moving cells reconcile these different dynamics. The RhoA-Rac1 interaction network model defines minimal, autonomous biochemical machinery that is necessary and sufficient for biologically observed modes of cell movement.

## Results

### Spatially variable topology of the RhoA-Rac1 interaction network

The Rac1 effector PAK inhibits RhoA, and the RhoA effector ROCK inhibits Rac1 ^16^. Here, we tested how the other RhoA effector, DIA, influences the Rac1 and RhoA activities. We first downregulated DIA using small interfering RNA (siRNA) and measured the resulting changes in the Rac1-GTP and RhoA-GTP levels. Downregulation of DIA increased the RhoA abundance and decreased Rac1 abundance, while decreasing relative activities of both RhoA and Rac1 Figures S1A and S1B). The decrease of relative Rac1 and RhoA activities induced by DIA knockdown shows that DIA activates Rac1 and also supports the existence of a positive feedback loop between DIA and RhoA described earlier ^27^. In addition, the GTPase network features another positive feedback from PAK to Rac1 through several molecular mechanisms ^28–31^. Summing up the interactions between RhoA and Rac1 mediated by their effectors ROCK and PAK ^17^ and RhoA - Rac1 interactions through DIA, we arrive at the intertwined negative and positive feedback circuitry of the RhoA-Rac1 network shown in Fig. S1C.

To explain the distinct GTPase activities at the leading and trailing edges, we hypothesized that these diverse feedforward and feedback mechanisms may be spatially controlled. Therefore, we explored how the interactions of active RhoA with its effectors vary spatially in polarized MDA-MB-231 cells. Using a proximity ligation assay (PLA), which visualizes protein interactions *in situ* ^32,33^, we measured RhoA-DIA and RhoA-ROCK complexes (Figure 1A and 1B). Based on the commonly considered morphology of the long, narrow cell rear and the wide leading edge ^34^, we segmented each polarized cell into three parts: the rear (about 20% of the cell length), intermediate region (next 70% of the cell length), and front (the rest 10% of the length). The density of the RhoA-effector complexes was quantified by dividing the number of PLA reactions by the area of the corresponding compartment.

**Figure 1.**
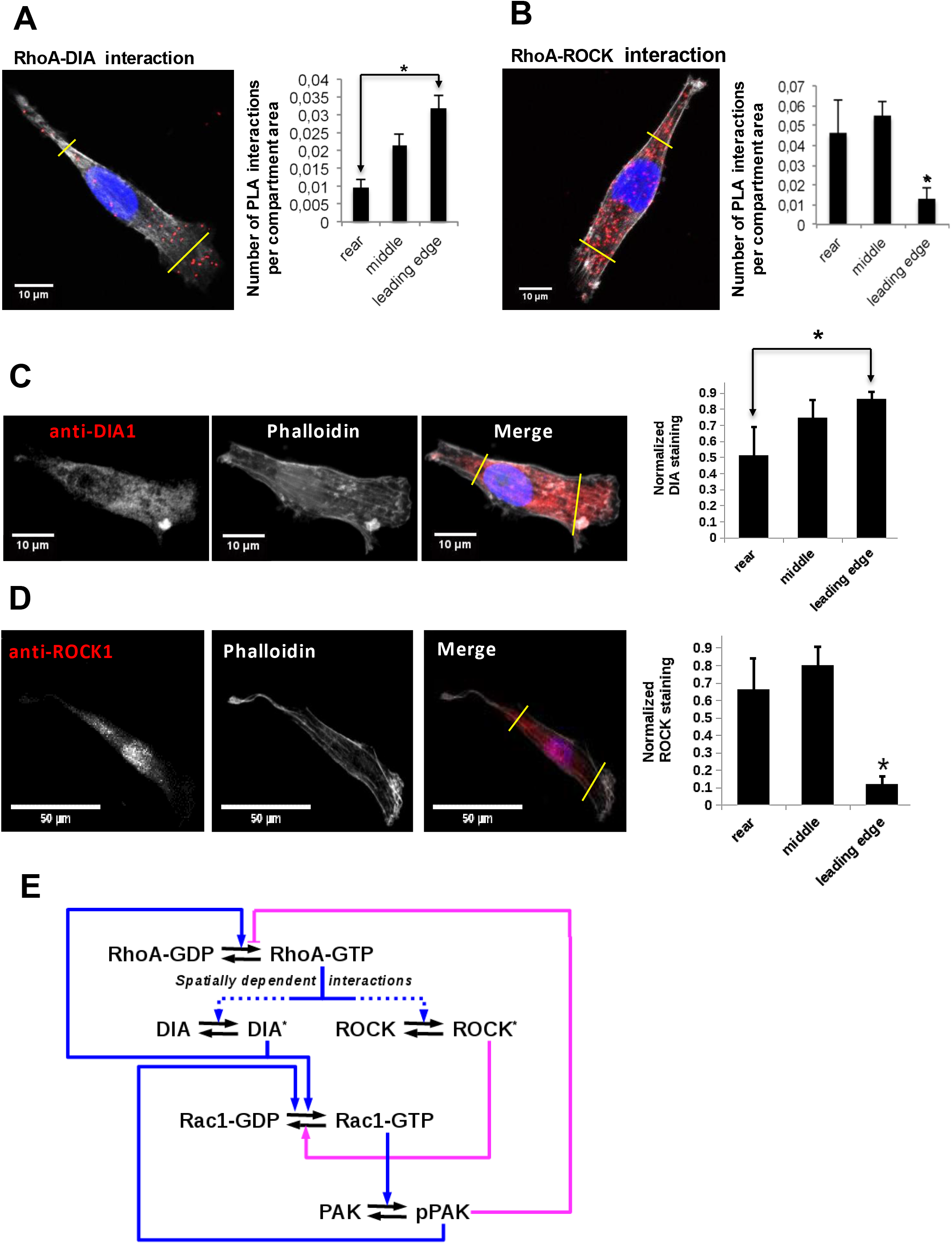
Differential localization of the RhoA-DIA and RhoA-ROCK1 protein complexes determine spatially resolved signaling topology. (**A, B**) Representative PLA images. Each red spot within a cell represents a fluorescent signal from a single RhoA-DIA1 (**A**) or RhoA-ROCK1 (**B**) complex. Yellow lines indicate bounds for the leading edge, intermediate region and rear. Bar graphs at the right show the average density of these complexes in different cell regions (the rear, middle and leading edge) ± S.E.M. of four independent experiments with 25 cells analyzed per experiment. The asterisk * indicates that p<0.05 calculated using unpaired t-test. (**C, D**) Representative images of DIA1 and ROCK1 immunostaining. Bar graphs at the right show quantified immunostaining density signals for different cellular compartments ± S.E.M. of four independent experiments with 1 cell analyzed per experiment. The asterisk * indicates that p<0.05 calculated using unpaired t-test. (**E**) A schematic wiring diagram of the RhoA-Rac1 network, showing positive (blue) and negative (magenta) feedback loops. Spatially varying RhoA interactions with its effectors DIA and ROCK are shown by dashed lines.

The results show that the RhoA-DIA complexes are predominantly localized at the cell front, whereas their density is markedly decreased at the rear (Figure 1A). In contrast, the density of the RhoA-ROCK complexes increases towards the cell rear and decreases at the leading edge (Figure 1B). These results are in line with protein staining data in polarized cells, which suggest that DIA is mainly localized at the leading edge (Fig. 1C), whereas ROCK is abundant at the rear and cell body (Fig. 1D) ^24,35–38^. For MDA-MB-231 cells, our quantitative proteomics data showed that the RhoA abundance is at least 10-fold larger than the abundance of DIA and ROCK isoforms combined ^17^. Thus, as shown in the Section 1 of Supplemental Methods, the RhoA-effector concentrations depend approximately linearly on the DIA and ROCK abundances. Taken together, these results suggested a protein interaction circuitry of the GTPase network, where competing effector interactions are spatially controlled (Fig. 1E). In order to analyze how this differential spatial arrangement of GTPase-effector interactions can accomplish the dynamic coordination between the leading and trailing edge, we constructed a mechanistic mathematical model and populated it by quantitative mass spectrometry data on protein abundances (Table S1).

### Analyzing the dynamics of the RhoA-Rac1 interaction network

The changes in ROCK and DIA abundances along the longitudinal axis of polarized cells (Fig. 1C,D) could plausibly encode the distinct RhoA-Rac1 temporal behaviors in different cellular segments. Therefore, we explored these possible dynamics of the GTPase network for different DIA and ROCK abundances prevailing at different spatial positions along the cell length. We first used a spatially localized, compartmentalized model where different DIA and ROCK abundances corresponded to distinct spatial locations (see Section 2 of Supplemental Methods for a detailed description of this model).

Using the model, we partitioned a plane of the ROCK and DIA abundances into the areas of different temporal dynamics of RhoA and Rac1 activities (Fig. 2A). This partitioning is a 2-parameter bifurcation diagram where the regions of distinct GTPase dynamics are separated by bifurcation boundaries at which abrupt, dramatic changes in the dynamic behavior occur. The blue region 1 in Fig. 2A corresponds to the self-perpetuating oscillations of the RhoA and Rac1 activities at the leading edge. There, the ROCK abundance is markedly lower and the DIA abundance is higher than in the cell body (Fig. 1C and 1D). Thus, at the leading edge a combination of Rac1 activation by RhoA via DIA and RhoA inhibition by Rac1 via PAK (Fig. 2B) results in sustained oscillations of RhoA and Rac1 activities (Fig. 2D). This periodic Rac1 activation drives actin polymerization at the leading edge pushing protrusion-retraction cycles ^6,18,25,39^.

**Figure 2.**
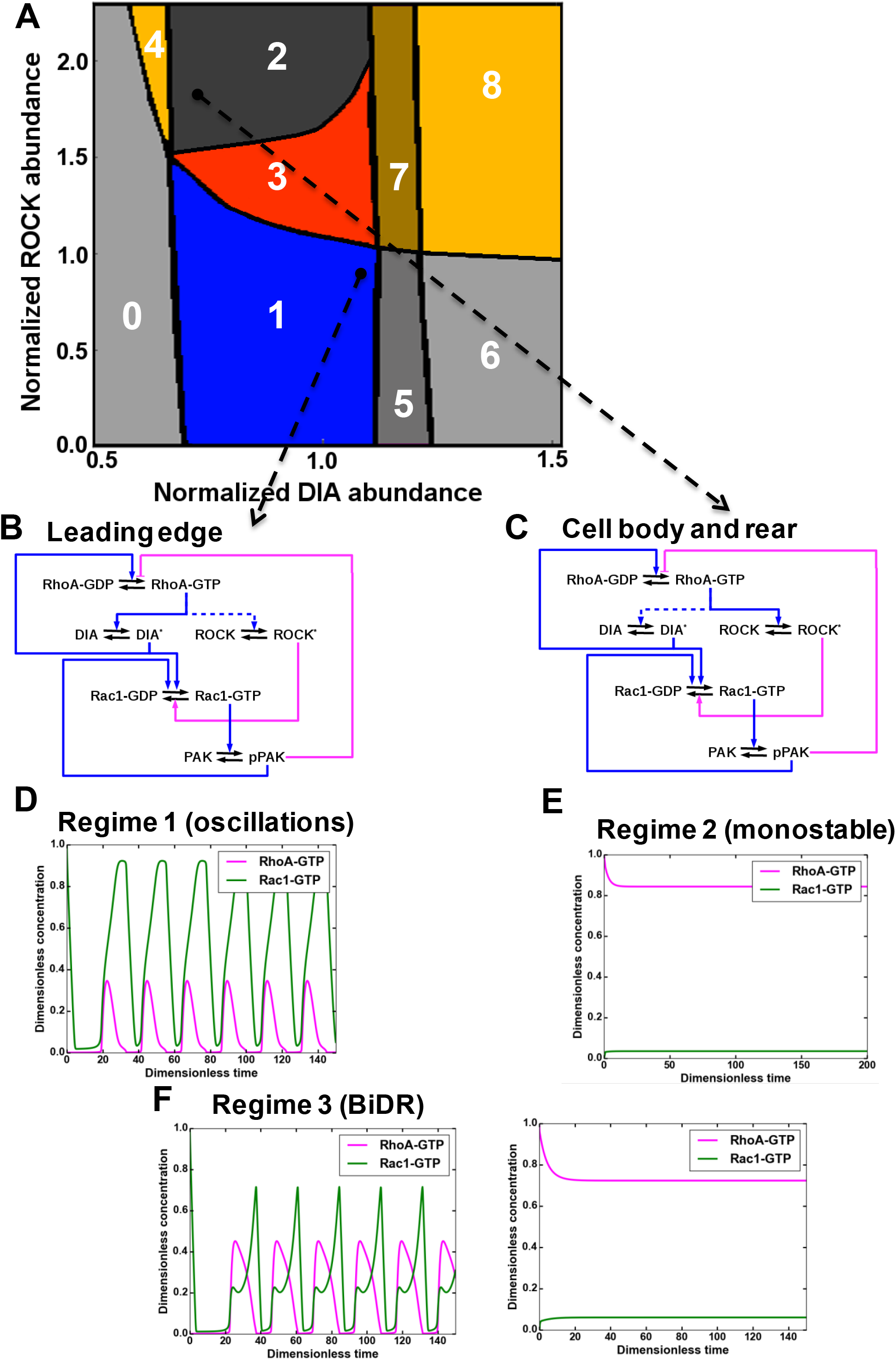
A mathematical model of the RhoA-Rac1 network predicts dramatically distinct dynamic regimes for different DIA and ROCK abundances. (**A**) Distinct dynamic regimes of the RhoA-Rac1 network dynamics for different DIA and ROCK abundances. Oscillations of RhoA and Rac1 activity exist within area 1 (regime 1). In area 3, sustained GTPase oscillations and a stable steady state with high RhoA and low Rac1 activities coexist. Regimes 0, 2, 5 and 6 have only one stable steady state. Steady state solutions with high RhoA activity exist in areas 2-4, and 6-8. Stable steady state solutions with high Rac1 activity exist in areas 0 and 5-8. Regimes 4, 7 and 8 are bistable with two stable steady states. (**B, C**) Wiring diagrams of the RhoA-Rac1 network for the cell leading edge (**B**) and the cell body and rear (**C**). Dashed blue lines indicate weak activating connections. (**D-F**) Typical time courses of RhoA and Rac1 activity in regimes 1 (**D**), and 2 (**E**). (**F**) In area 3, depending on the initial state, the GTPase network evolves either to a stable steady state (right) or a stable oscillatory regime (left).

The dark grey region 2 in Fig. 2A is an area of stable high RhoA and low Rac1 activities at the rear and intermediate cell regions. Within these regions, RhoA inhibits Rac1 via ROCK, and Rac1 inhibits RhoA via PAK (Fig. 2C), and after perturbations, the GTPase network converges to steady-state levels of high RhoA-GTP and particularly low Rac1-GTP (Fig. 2E).

The red region 3 corresponds to the coexistence of GTPase oscillations and a stable steady state with high RhoA and low Rac1 activities. Depending on the initial state, the GTPase network evolves to different dynamic regimes. If the initial state has high RhoA-GTP and low Rac1-GTP, the GTPase network progresses to a stable steady state, but if the initial state has low RhoA-GTP and high Rac1-GTP, the network will develop sustained oscillations (Fig. 2F). This region 3 is termed a BiDR (<u>B</u>i-<u>D</u>ynamic-<u>R</u>egimes) by analogy with a bi-stable region where two stable steady states coexist and the system can evolve to any of these states depending on the initial state ^23^. However, in contrast with bistable regimes only one of two stable regimes is a stable steady state in the BiDR region, whereas the other dynamic regime is a limit cycle that generates stable oscillations.

In addition to these dynamic regimes, the spatially localized model predicts other emergent non-linear dynamic behaviors (Fig. 2A, Figs. S2A-D and Fig. S3), which the GTPase network may execute under large perturbations of the RhoA and Rac1 effector abundances to coordinate GTPase signaling at the leading and trailing edges (see Section 2 of Supplemental Methods for a detail description of these regimes). Therefore, we next analyzed how the leading and trailing edge GTPase dynamics are coupled.

### Spatiotemporal dynamics of the RhoA-Rac1 network reconciles the distinct temporal behaviors at the cell front and rear

Different active GTPase concentrations in the cell rear and the leading edge induce diffusion fluxes ^40^, which in turn influence the emerging behavior of these GTPases and coordinate their dynamics in distinct cellular segments. Therefore, we first explored the behavior of the RhoA-Rac1 network in space and time using a spatiotemporal model of the GTPase network interactions (referred to as a reaction-diffusion model, see Section 2 of Supplemental Methods). We digitized 2D images of polarized cells and incorporated the DIA and ROCK abundances as functions of the spatial coordinate along the cell length, based on the quantitative imaging data (Fig. 3A, B, C).

**Figure 3.**
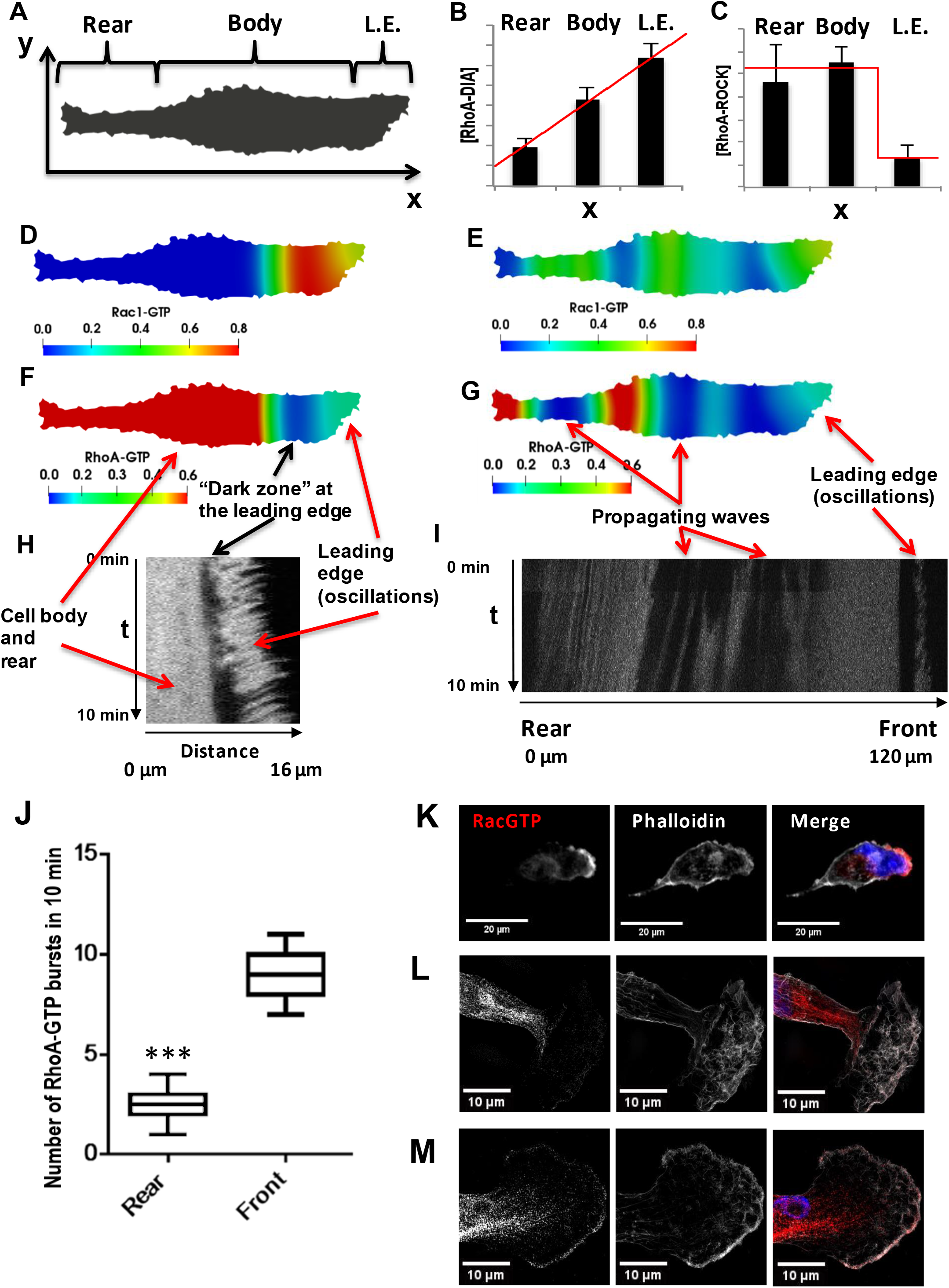
Spatial propagation of RhoA and Rac1 activities during cell motility. (**A**) A 2-D calculation domain obtained by digitizing cell images. Different cellular compartments are indicated. The x-axis represents the direction of cell polarization, the y-axis represents the perpendicular direction. (**B, C**) The abundance profiles of DIA and ROCK used in simulations (red lines) are superimposed on the experimental spatial profiles (bar graphs in Figs. 1C,D). (**D-G**) Model-predicted spatial patterns of the RhoA and Rac1 activities for different phases of the cell movement cycle. (**D, F**) Rac1 and RhoA activity snapshots during a protrusion-retraction cycle at the leading edge. (**E, G**) represent snapshots when the Rac1 and RhoA activity wave have spread over the entire cell, reaching the rear. (**H**) The RhoA activity at the leading edge and cell body during a protrusion-retraction phase measured by RhoA FRET probe in space and time. The arrows compare model-predicted and experimentally measured patterns, indicating zones of RhoA oscillatory and high constant activities and a “dark zone” of low RhoA activity. (**I**) Spatiotemporal pattern of the RhoA activity during further RhoA wave propagation into the cell. (**J**) The number of RhoA activity bursts at the cell body and rear during 10 minutes measured using the RhoA FRET probe. Error bars represent 1^st^ and 3^rd^ quartiles, *** indicate p < 0.001 calculated using unpaired t-test. (**K-M**) Fluorescent microscopy images of Rac1 activity (red), combined with staining for F-actin (phalloidin, white) and the nucleus (DAPI, blue) in fixed cells for different phases of the cell movement cycle; (**K**) a protrusion-retraction cycle at the leading edge, and (**L, M**) present Rac1 activity wave propagation into the cell body. The images (**L, M**) were obtained by super-resolution microscopy.

The model predicts autonomous, repeating cycles of the spatiotemporal GTPase dynamics (Figures 3D-G and Supplemental Video S1). For a substantial part of a dynamic cycle, high RhoA-GTP and low Rac1-GTP persist at the cell rear and maintain the rear contraction, whereas active RhoA and Rac1 oscillate at the leading edge, resulting in actin (de)polarization cycles and protrusion-retraction cycles (Figs. 3D and 3F) ^9^. At the same time a wave of oscillating Rac1 and RhoA activities slowly propagates from the leading edge towards the cell rear (Figs. 3E and 3G). Between the oscillatory RhoA-GTP zone and the areas of high RhoA activity, a zone of low RhoA activity emerges (Fig. 3F). As time progresses, the wave of oscillating GTPase activities and the area of low RhoA activity spread to the rear (Figs. S4A and S4B), leading to re-arrangement of the cytoskeleton and dissociation of focal adhesions. Because of the oscillations, zones of low Rac1 activities emerge, which give rise to high RhoA-GTP that interacts with ROCK and leads to the rear retraction (Supplemental Video S1). Subsequently, RhoA returns to its initial high stable activity, and the dynamic pattern of RhoA-GTP and Rac1-GTP over the entire cell returns to its initial state.

To test the model predictions, we used cells stably expressing the mTFP-YFP RhoA-GTP FRET-probe ^41^ allowing us to determine the RhoA-GTP dynamics using ratiometric, live-cell spinning disk microscopy. We imaged the cells with a frequency of one image every five seconds and constrained the measurement time to 10 minutes to limit phototoxic effects. Due to this time limitation, a full cycle of cellular movement (around 45 minutes on average, Supplemental Video S2) could not be followed in an individual cell, and the full spatiotemporal RhoA activity cycle during a cell movement was compiled from several cells observed in different phases of cellular movement. In the initial part of the cell movement cycle, the spatiotemporal RhoA activity showed three different zones: (*i*) oscillations at the leading edge, (*ii*) dark zone of low activity and (*iii*) light zone of high activity (Figs. 3H and S4C) in the cell body and rear, matching the model prediction (Fig. 3F). As time progressed, the GTPase activity wave propagated further into the cell (Fig. 3I), forming zones of high and low RhoA activities. In the space-time coordinates, the slope of the boundaries of these zones suggests that they travel from the leading edge to the cell rear, confirming the model predictions (cf. Supplemental Video S1 and Fig. 3I). When the wave of oscillatory GTPase activities finally reaches the cell rear, it induces several RhoA-GTP spikes (Figs. 3G and 3I), periods of low RhoA activity (Figs. S4A-B and S4D), and subsequent return to the original, high RhoA-GTP at the rear and part of the cell body (Figs. 3F and 3H). Fig. S4D experimentally captures this transition from a low RhoA activity to the original high activity as the final step of the cell movement cycle predicted by the model.

The model predicts that during a single cellular movement cycle, multiple bursts of RhoA activity appear at the leading edge, whereas at the cell rear, RhoA activity bursts occur only after the RhoA-Rac1 wave has spread through the cell (Supplemental Video S1). Measuring the number of RhoA bursts at the leading edge and cell rear during observation time (10 min) corroborated model predictions, showing a ca. 5-fold larger number of bursts at the leading edge than at the cell rear (Fig. 3J). On average, at the leading edge a burst of RhoA activity happens every minute, while at the cell rear only 1 or 2 bursts happen during 10 minutes (Fig. 3J).

Although spatially resolved Rac1 activity can be determined using exogenous probes, they dramatically change the cell shape when expressed ^18^. However, endogenous Rac1-GTP can be reliably detected by immunostaining with a conformation-specific Rac1-GTP antibody. Rac1 was mainly active at the leading edge with lower activity in the space between the nucleus and cell rear (Figure 3K), similar to the patterns observed in the model for protrusion-retraction cycles (Fig. 3D). The GTPase waves can be detected using super-resolution imaging. These images corroborated the Rac1-GTP presence towards the cell nucleus and rear (see super-resolution images in Figs. 3L-M and S4E). The series of images shown in Figs. 3K-M and S4E is consistent with the concept of traveling Rac1-GTP waves predicted by the model.

These spatiotemporal activation dynamics of Rac1 and RhoA underlie the morphological events during cell migration, i.e. protrusion-retraction cycles at the front and the adhesion-retraction cycle at the rear ^1^ (Supplemental Video S2). In the initial phase of cell migration the Rac1-RhoA oscillations push out and retract lamellopodia at the leading edge permitting the cell to explore its environment and follow chemotactic cues ^6^, while high RhoA activity at the trailing edge stabilizes cell adhesion ^42^. In the late migration phase, RhoA activity extends towards the front allowing focal adhesions to form at the front, and stress fibers to generate contractile force in the cell body that will retract the rear. At the same time, Rac1 activity traveling towards the trailing edge destabilizes focal adhesions at the rear. The combination of these activities pulls up the rear resulting in cell movement. Their critical coordination is accomplished by the intertwined spatiotemporal dynamic regulation of Rac1 and RhoA described by our mathematical model.

### Hysteresis of Rac1 and RhoA activities and cell shape features

We previously showed that PAK inhibition could change the cell shape of MDA-MB-231 cells from mesenchymal to amoeboid ^17^. The mesenchymal mode of migration features an elongated cell morphology and high Rac1 activity, whereas the amoeboid mode is hallmarked by a rounded morphology and high RhoA activity ^21^. These morphologies and migration types are mutually exclusive but can transition into each other. Our previous study showed that this transition correlated with the hysteresis of active RhoA and Rac1 upon PAK inhibition ^17^. Hysteresis is the hallmark of bistability: if a parameter, such as the PAK abundance, reaches a threshold value, then the system flips from one stable state to another stable state, at which it remains for a prolonged period of time even when this parameter has returned to its initial value.

Our model allows us to examine the exact spatiotemporal kinetics of the GTPase network in response to changes in PAK abundance or activity, which causes Rac1 and RhoA activities to move through different dynamic regimes (shown by the line connecting points I – III in Fig. 4A). In unperturbed cells, GTPase activities oscillate at the leading edge. This initial network state corresponds to point I in region 1 and unperturbed ROCK, PAK and DIA abundances and activities (Fig. 4A). Because Rac1 and RhoA are difficult to target for therapeutic interventions, we used a small molecule PAK inhibitor (IPA-3) in our previous study ^17^. As the PAK abundance gradually decreases (or PAK inhibition increases), the system moves from the oscillatory region 1 to the BiDR region 3 before it reaches a bistable regime (regions 7 and 8), as shown by point II. In the BiDR region a stable high RhoA-GTP, low Rac1-GTP state and a stable oscillatory state with a high average Rac1-GTP coexist at the leading edge (Fig. 2F). In the bistable regimes 7 and 8, two stable states co-exist (*i*) high RhoA-GTP, low Rac1-GTP; and (*ii*) low RhoA-GTP, high Rac1-GTP (Fig. S3). Plotting the trajectory of the changing network states in response to a decrease in PAK abundance (blue curves in Figs. 4B and 4C) shows that Rac1 activity (averaged over cell volume and time for the oscillatory dynamics) first gradually decreases and then abruptly decays after passing point II (Fig. 4B). If we follow the Rac1-GTP trajectory in response to increasing PAK inhibitor doses, we obtain a similar curve (Fig S5). The average RhoA-GTP behaves oppositely, steadily increasing and then jumping to peak activity after the network passes the BiDR and bistable regions (blue curves in Fig. 4C and S5B showing the RhoA-GTP trajectories in response to PAK abundance decrease or IPA-3 increase, respectively). A further decrease in the PAK abundance moves the RhoA-Rac1 network into point III of region 6 with a single steady state of active RhoA and low Rac1 activity (Figs. 4A-C).

**Figure 4.**
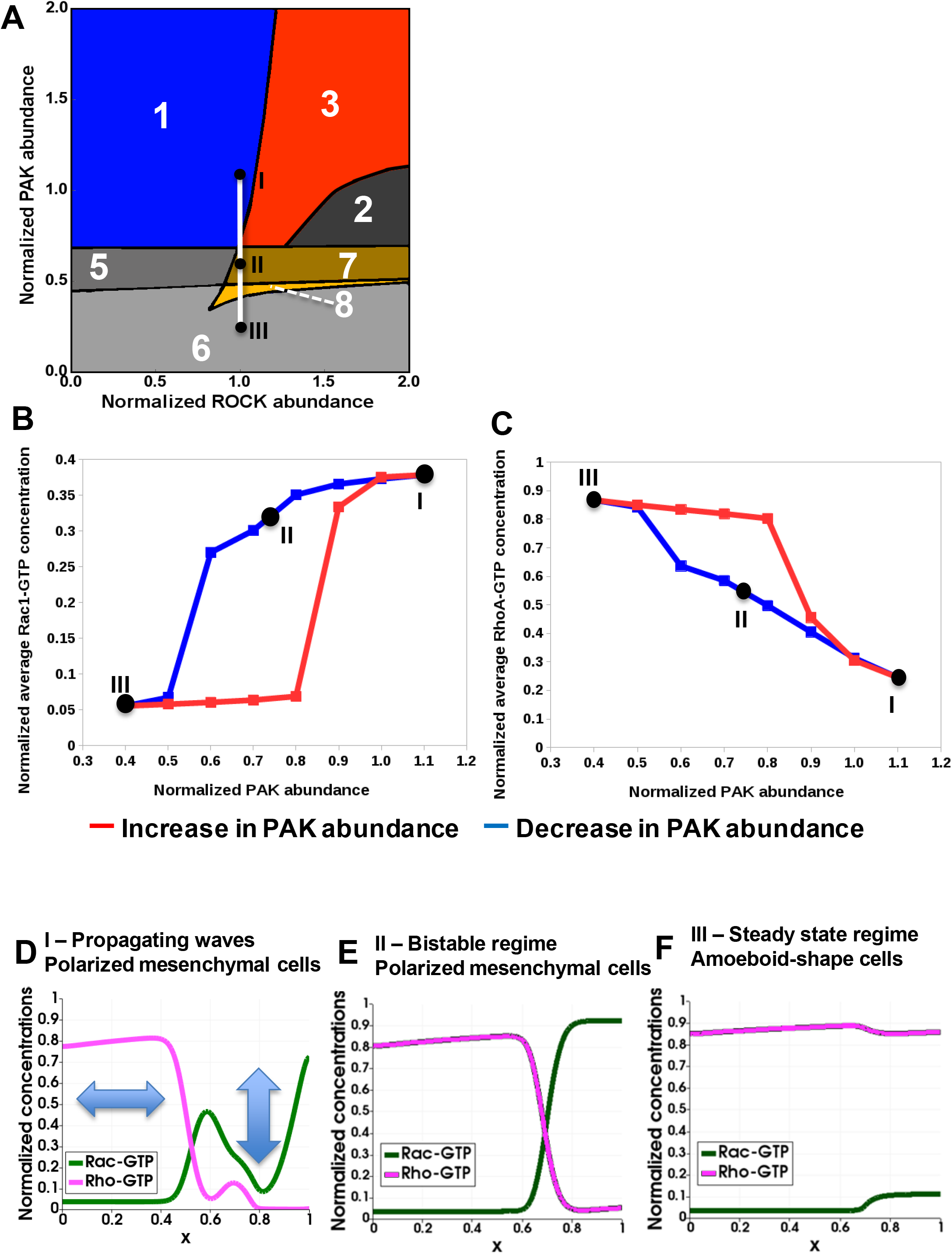
Hysteresis of the RhoA and Rac1 activities are manifested upon PAK inhibition and recapitulated by a spatiotemporal model. (**A**) Distinct dynamic regimes of the RhoA-Rac1 network for different DIA and ROCK abundances. Colors and numbers of dynamic regimes are the same as in Fig. 2A. (**B, C**) Model-predicted dependencies of the RhoA and Rac1 activities on the PAK abundance for gradually decreasing (blue) and increasing (red) PAK abundances. The network evolution occurs through two different routes (blue and red curves in Figs. 4B and 4C). It is calculated by averaging the GTPase activities over the time and cell volume based on Western blot data reported in our previous study ^17^. Points I, II and III shown in black (**A**) are also indicated on the network trajectories (**B, C**). (**D-F**) Snapshots of simulated RhoA-GTP and Rac1-GTP spatiotemporal patterns that emerge for different PAK abundances are shown for a 1-D section of a cell. The x axis corresponds to the normalized cell length (Fig. 3A). Arrows in panel (**D**) illustrate oscillations and the wave propagation along a cell.

The spatiotemporal dynamic pattern corresponding to point I (Figs. 4B and 4C) is a propagating wave illustrated in Fig. 3D-H and schematically shown in Fig. 4D where the arrows illustrate oscillations and the wave propagation along a cell. For point II, the RhoA and Rac1 activity patterns depend on space, but do not change with time (Fig. 4E). Such spatial dynamics are referred to as a pinning or stalled wave, meaning that a wave of activation first propagates in space, then decelerates and eventually stops, forming stationary RhoA and Rac1 activity profiles ^14^ with high steady-state Rac1-GTP at the leading edge (Fig. 4E). Phenotypically cells maintain a mesenchymal state and a polarized shape in both states I and II (Fig. 3I and 3J). For point III the resulting steady-state profile features high RhoA and low Rac1 activities along the entire cell (Fig. 4F), which is a hallmark of amoeboid cells ^21,43,44^. Our results suggest that the transition from the mesenchymal to the amoeboid phenotype becomes switch-like once PAK activity falls below a critical threshold ^17^.

What about the transition back? Because the underlying GTPase activities show hysteretic behavior, the transition from amoeboid back to the mesenchymal state should follow a different path. Indeed, in our previous study we observed that the switch from the mesenchymal to amoeboid state occurred at a higher level of PAK inhibition than the switch back when inhibition was gradually reduced ^17^. Our model now can explain the underlying spatiotemporal GTPase dynamics. If cells are forced into the amoeboid state by inhibiting PAK and then allowed to gradually regain PAK activity (red curves in Figs. 4B, C and S5), the network does not pass through the stalled wave state (point II in Figs. 4B and C). It rather first moves from point III in region 6 through bistable regions maintaining high RhoA and low Rac1 activities. Upon further relief of PAK inhibition, the network then passes through the BiDR region, and the Rac1 activity jumps to a high value, whereas the RhoA activity switches to a low value, approaching initial point I (Figs. 4B,C).

Summarizing, the experimentally observed hysteresis of RhoA and Rac1 activities upon PAK inhibition is explained by the network evolution through the BiDR and bistable regions. The morphological cell shape changes also follow this pattern. Importantly, bistability in the RhoA-Rac1 network only can be achieved through PAK inhibition, and only when PAK is largely inhibited, cells leave the bistable regions and reach a stable state III where their cell shape becomes amoeboid ^45^.

### ROCK inhibition results in multiple competing lamellipodia and multi-polar cell shapes

Having investigated the consequences of PAK inhibition, we next studied the effects of ROCK inhibition. The model predicts that a decrease in ROCK activity below a certain threshold results in the formation of several oscillatory centers of GTPase activities featuring high (averaged over time) Rac1 activity (Supplemental Video S3). In contrast to periodic GTPase waves propagating from a single Rac1 oscillatory center at the leading edge, several oscillatory Rac1 activity centers result in the uncoordinated and chaotic emergence of waves, thereby preventing a single wave propagation along a cell (compare Supplemental Videos S1 and S3). These findings might imply the emergence of multi-polar cells that extend lamellipodia in several different directions. In fact, multiple competing lamellipodia emerging as a result of ROCK inhibition were previously reported ^46^.

To determine if ROCK inhibition could induce multiple Rac1-GTP foci, we seeded MDA-MB-231 cells on collagen and treated the cells with the pan-ROCK inhibitor Y-27632. After 15 minutes we fixed the cells and stained for active Rac1 and F-actin. Spatially resolved Rac1 activity showed two or three Rac1-GTP poles, whereas cells not incubated with the inhibitor were exclusively mono-polar (Figures 5A and S6A). Measured using the RhoA-GTP FRET-probe, patterns of the RhoA activity (Fig. 5B) showed chaotic spontaneous activity bursts, as well as periods of relatively steady, high-RhoA activity in the central part of the cell. These dynamics are in line with model-predicted patterns (Supplemental Video S3), and in a sharp contrast to cells where ROCK is not inhibited (Fig. 3).

**Figure 5.**
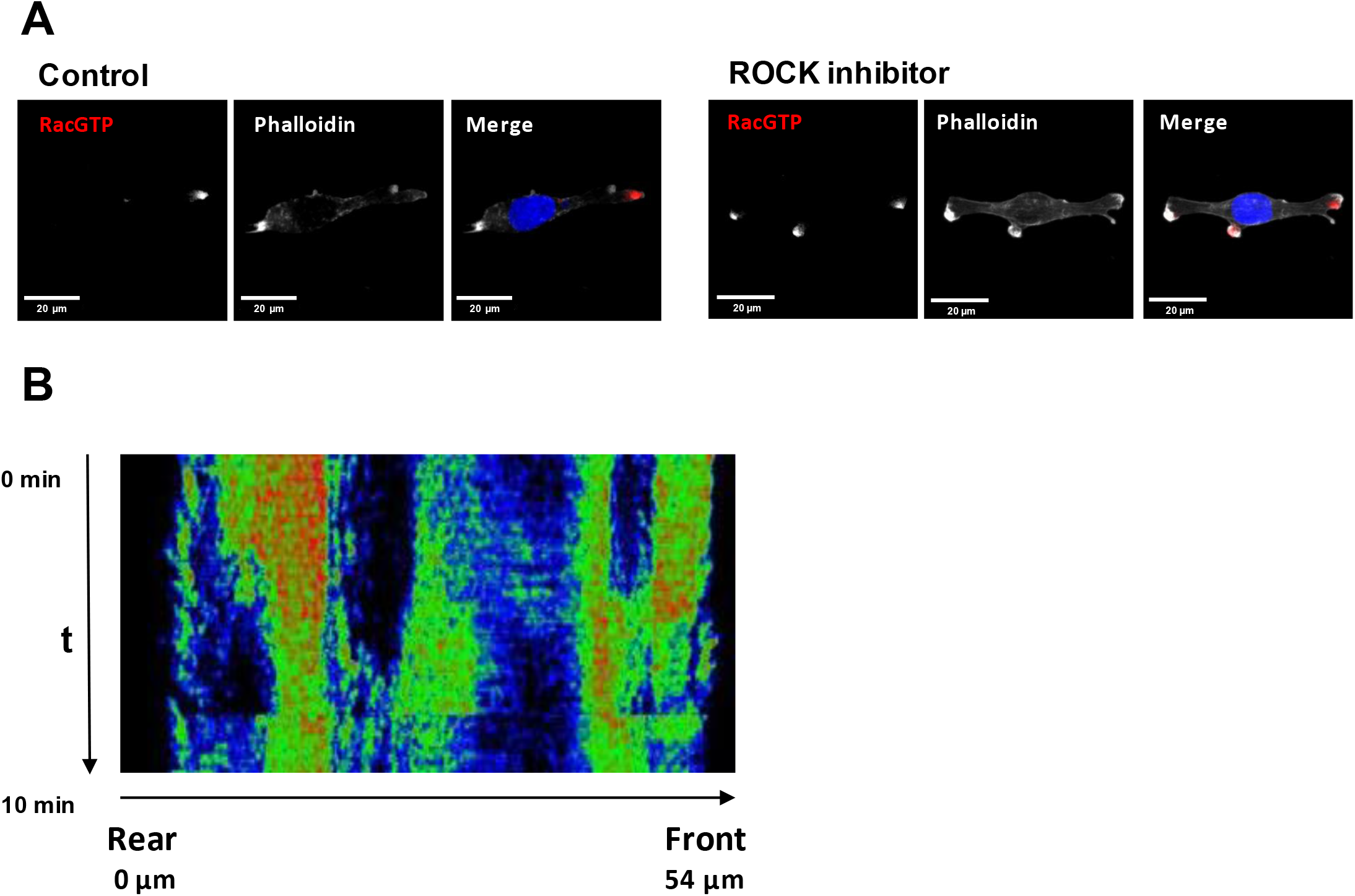
Inhibition of ROCK leads to the formation of multi-polar cells. (**A**) Fluorescent microscopy images of Rac1 activity (red), and F-actin (phalloidin, white) and nuclear (DAPI, blue) staining in fixed MDA-MB-213 cells treated or not with 2.5μM Y-27632 ROCK inhibitor for 15 min. (**B**) Spatiotemporal pattern of the RhoA activity in cells treated with 2.5μM of ROCK inhibitor Y-27632 measured by the RhoA FRET probe.

In the absence of ROCK inhibitor, the RhoA-GTP bursts at the cell rear only occur when a propagating wave reaches the rear, i.e. at low frequency. These bursts cause the cell tail retraction and are associated with the last step of the movement cycle of a polarized elongated cell. When ROCK is inhibited, a GTPase oscillatory center emerges in the tail with the corresponding increase in the frequency of RhoA-GTP bursts (Figs. S6B, C). As a result, a cell loses the ability to retract the tail. These cells do not lose polarity but exhibit substantial morphological changes, acquiring largely elongated shapes (compare Supplemental Videos S2 and S4). In line with these results, our experiments suggest that the total migration distance is smaller for cells treated with ROCK inhibitor than for untreated cells (Fig. S6D). This decrease can be explained by the formation of multiple lamellipodia and the inability of ROCK-inhibited cells to retract their tail.

Summing up, these data suggest that the ROCK activity above a certain threshold is necessary for the formation of a single high Rac1 activity center at the leading edge and avoiding the appearance of multiple high Rac1 activity centers in a cell. Thus, ROCK cooperates with PAK to maintain the polarized lamellipodia formation and the cell shape typical for mesenchymal cell movement.

## Discussion

Rho GTPases are core regulators of mesenchymal and amoeboid cell migration. They integrate multiple internal and external cues ^47–51^ and relay information to a variety of cellular protein machineries, including proteins driving actin polymerization and cytoskeleton rearrangements, thereby enabling cell migration^52^. Although molecular details of Rho GTPase - effector interactions have been elaborated, we still lack an overall picture of how these GTPase activities and effector interactions are coordinated between the leading and trailing edge in order to enable cell movement. Here we present a minimal biochemical mechanism that is necessary and sufficient for the cyclic process of cell migration. This mechanism integrates different temporal dynamics of the RhoA and Rac1 GTPases at the cell front, body and rear and shows how these activities are coordinated by propagating GTPase activation waves. Moreover, our model can rationalize how the amoeboid and mesenchymal types of migration interchange by suppression or over-activation of specific RhoA and Rac1 effectors.

A traditional view on mesenchymal migration is that high Rac1 activity persists only at the leading edge, whereas high RhoA activity exists mainly at the rear. This view is supported by the reported mutual antagonism of Rac1 and RhoA ^17,21^. However, live cell imaging experiments showed oscillations in RhoA activity at the leading edge, challenging the traditional view ^6,18,25^. Several studies suggested that RhoA not only inhibits Rac1 via ROCK but also activates Rac1 via DIA ^16,53^. Our results and literature data show the differential spatial localization of DIA and ROCK within the cell (Figs. 1C and 1D, ^24,35–38^). ROCK is more abundant at the cell rear and body, whereas DIA is more abundant at the leading edge. This leads to marked changes in the cellular distribution of RhoA-ROCK versus RhoA-DIA effector complexes (Figs. 1A and 1B). Differential localization of DIA and ROCK (as well as different spatial distribution of GEFs, GAPs, and guanosine nucleotide dissociation inhibitors ^26,54,55^) can generate distinct circuitries of RhoA-Rac1 interactions and different RhoA and Rac1 kinetics along a cell (Fig. 2B-F). Oscillations of RhoA and Rac1 activities at the leading edge guide protrusions and retractions, whereas high, stable RhoA activity and low Rac1-GTP at the rear maintain focal adhesions that assure cell attachment to the substrate (schematically illustrated in Figures 2B). Although the distinct Rho GTPase dynamics at the front and rear during a cell migration cycle have been described, it is unknown how exactly a cell integrates these behaviors to coordinate cell movement.

To better understand these complex kinetics we have developed a model of the GTPase dynamic behaviors in time and space. Our model suggests that periodically repeating GTPase waves connect protrusion-retraction oscillations of the GTPase activities at the leading edge and almost stable RhoA and Rac1 activities at the rear. These waves occur due to diffusion fluxes that are induced by different RhoA-GTP and Rac1-GTP concentrations along the cell and the excitable dynamics of RhoA and Rac1 generated by negative and positive feedback loops in the network ^26^. When a wave that propagates from the front reaches the rear, the emerging Rac1-GTP spikes can induce the dissociation of focal adhesions and cytoskeleton reassembly, while intermittent RhoA-GTP spikes of high amplitude (Supplemental Video S1) can force the cell rear retraction and movement. Subsequently, the RhoA and Rac1 activities at the rear return to their stable levels. These RhoA and Rac1 activity waves create an autonomous, cyclic mechanism that controls the mesenchymal type of cell migration.

Model predictions are supported by imaging and Western blot experiments. Experiments with the RhoA FRET probe corroborated the predictions of RhoA-GTP dynamics at the leading edge (Fig. 3H) and cell body and rear (Figs. 3I, 3J and S4C). Cell staining with specific Rac1-GTP antibody provided snapshots of Rac1 activity corresponding to protrusion-retraction cycles (Fig. 3K) and the spreading of Rac1 activity beyond the leading edge into the cell body (Figs. 3L, 3M and S4C, super-resolution microscopy images) as predicted by the model.

Whereas PAK inhibition (Fig. 4) induces a transition from the mesenchymal to amoeboid mode of migration and the corresponding changes in the cell shapes ^17^, ROCK inhibition can lead to the formation of multiple centers of Rac1 oscillations (Figure 5) and multiple competing lamellipodia ^46^. At the same time, DIA downregulation by siRNA resulted in substantial rewiring of the RhoA-Rac1 signaling network, manifested by an increase in RhoA abundance and a decrease in Rac1 abundance (Figs. S1A and S1B). Model simulations show that these changes can decrease a threshold DIA abundance required to maintain the initial GTPase dynamics in time and space (Fig. S6E). Thus, cells tend to adapt to DIA1 perturbation by adjusting other protein abundances to keep a minimally perturbed Rho-Rac signaling pattern.

Actin travelling waves arising from signal transduction excitable network (STEN) were previously described, and elegant mathematical models were proposed that analyzed the dynamics of various small networks of cytoskeleton proteins and GTPases ^51,56–58^. These models, together with a more abstract model considering generic activators and inhibitors ^59^, explained the observed wave-like signal transduction patterns and actin waves, which were localized to the cell front, driving protrusion-retraction cycles ^60^. The periodic waves of Rac1-RhoA activities described in this paper propagate through the entire cell, coordinating protrusion-retraction cycles at the front and the adhesion-retraction cycle at the rear, and therefore are very different from travelling waves reported previously.

In addition to diffusion and excitable properties of signaling networks, the cell front and rear can communicate via other molecular mechanisms. It was suggested that microtubules can play an important role in the spatial localization of GTPase related proteins and the coordination of front and back signaling ^61–63^. Staining intensities of F-actin at the front and phosphorylated myosin light chain 2 (pMLC2) at the rear showed that they were neither positively correlated nor anticorrelated ^9^. These readouts could be interpreted as the GTPase signaling activities at the front and the rear. The discovered buffering of the front and rear signaling was completely destroyed by the disruption of microtubules ^9^. Different spatial concentration profiles of RhoA and Rac1 downstream effectors considered in our model could conceivably depend on the microtubule network.

In summary, our spatiotemporal model of RhoA-Rac1 signaling proposes how different GTPase dynamics at the cell front and rear are coupled and explains the changes in signaling patterns and cell shapes upon inhibition of GTPase effectors. It represents a minimal, experimentally validated model of the biochemical GTPase network that regulates cell migration.

## Methods

### Experiments

#### Tissue Culture & cell treatment

##### Cells

MDA-MB-231 breast cancer cells (a gift from Brad Ozanne, Beatson Institute) were cultured in DMEM supplemented with 2 mM glutamine and 10% fetal calf serum at 37 °C in a humidified atmosphere containing 5% CO_2_. MDA-MB-231 expressing the RhoA activity probe were generated by lentiviral infection of the mTFP-YFP RhoA activity probe ^64^ and selected with puromycin at 2 μg/ml for three days. MDA-MB-231 cells with constitutive expression of nuclear mKATE2 were generated by infecting MDA-MB-231 cells with IncuCyte® NucLight Red Lentivirus Reagent (Cat. No. 4625) in the presence of polybrene (6μg/ml, Sigma). After 48h, selection was performed by supplement the media with puromycin (2μg/ml, Sigma).

##### ROCK inhibition

Cells were incubated with either vehicle, 1μM GSK 269962 (Tocris) or 2.5, 5 or 10μM (as indicated in manuscript) Y-27632 (Sigma) for 20min before the experiments were carried out.

##### Knock down by siRNA

Knock-down of DIA1 was achieved by transfecting a smartpool of three siRNAs targeting the human DIAPH1 mRNA and non-targeting siRNA control (Dharmacon cat. L-010347-00-0010). Both siRNAs were transfected at a final concentration of 50nM using Lipofectamine RNAiMax (Cat.13778) in a 1:2 (v/ v) ratio. Cells were kept for 48h before the experiments were carried out.

##### Rac1 and RhoA pulldowns

MDA-MB-231 and MDA-MB-231 transfected with siRNAs against DIAPH1 were seeded in a 6-well plate coated with rat-tail collagen (see siRNA experiments section) and lysed in 500 μl ice-cold lysis buffer (50 mM Tris-HCl, pH 7.5, 0.2% (v/v) Triton X-100, 150 mM NaCl, 10 mM MgCl_2_) supplemented with 1mM protease inhibitors PMSF and leupeptin (Sigma). Cell lysates were cleared of debris by centrifugation for 10 minutes at 20,000xg at 4°C. 10μl of the cleared lysate were kept as loading control. The remainder of the lysates were incubated with 6μl of GST-PAK-CRIB beads for Rac1 pulldowns or GST-Rhotekin-RBD beads for RhoA pulldowns for 1h at 4°C under end-to-end rotation. The GST-PAK-CRIB and GST-Rhotekin-RBD beads were produced as described by ^65^. The beads were washed with one volume of lysis buffer. The beads and an aliquot of the total lysate as input control were separated by SDS gel electrophoresis using 4-12% NuPAGE precast gels according to the manufacturer’s instructions. Gels were electroblotted onto PVDF membranes (Sartorius). Blots were blocked in TBST (50 mM Tris, pH 7.5, 150 mM NaCl, 0.05% Tween-20) containing 5% milk powder and incubated overnight with primary antibody followed by secondary antibodies linked to horse radish peroxidase (HRP). Antibodies used included: Rac1 antibody (Millipore, clone 238A, 1:500), anti-RhoA antibody (Santa Cruz Biotechnology 26C4, sc-418, 1μg/ml), anti-GAPDH (CST D16H11 XP®, diluted 1:3000) and anti-DIA1 (Thermo Fisher cat.PA5-21409, 1μg/ml). Secondary anti-rabbit and anti-mouse HRP-conjugated antibodies were obtained from CST and used at 1:10000 dilution. Western Blots were developed using SuperSignal™ West Femto Maximum Sensitivity Substrate (Thermo Fisher).

Images of the blots were acquired in a Bio-Rad ChemiDoc™ Imager. The Western blot bands were quantified using ImageJ.

##### Immunofluorescence

Cells were seeded onto high performance glass coverslip, thickness 1 1/2 (Zeiss, cat.474030-9000-000) coated with 0.01% collagen. For ROCK inhibition cells were pretreated as indicated in the corresponding section with Y-27632. Cells were washed twice with PBS, fixed and permeabilized with 3.7% formaldehyde, 0.025% NP-40 in 50mM Pipes pH6.8, 10mM MgCl_2_ for 5min. and blocked in TBS (50 mM Tris, pH 7.5, 150 mM NaCl) containing 2% BSA for 1h. Coverslips were incubated overnight in TBS containing 1% BSA with primary anti-Rac1-GTP (New East Bio cat.26903) (1:100), anti-ROCK1 (Thermo cat.PA5-22262) (1:100) or anti-DIA1 (1:200) antibodies. Slides were washed twice with TBS and then incubated for 1 h at room temperature with secondary antibodies anti-mouse F(ab’)2 Fragment Alexa Fluor ® 647 Conjugate (Thermo cat. A-21237), Donkey anti-rabbit Alexa fluor-488 (Cat. A-21206) or anti-rabbit Alexa fluor-594 (Thermo cat. A-11012) for confocal; anti-rabbit F(ab’)2 Fragment Alexa Fluor ® 594 Conjugate (Thermo cat. A-11072) and anti-mouse F(ab’)2 Fragment Alexa Fluor ® 647 Conjugate for super-resolution microscopy. Slides were washed twice with TBS and incubated with DAPI, 1:100, and phalloidin, conjugated with rhodamine or Alexa Fluor-488 (1:100) (Thermo A12379) for 5 minutes, washed two times and mounted using VECTASHIELD antifade mounting media (Vector labs Cat. H-1000). Confocal images were taken with an Olympus FV100 or a Nikon A1+ confocal, with 60x oil objective. Super-resolution images were taken with a N-SIM microscope using a with 100x oil objective.

##### Proximity ligation assay

The Proximity Ligation Assay (PLA) visualizes an interaction between two proteins that co-localize within ≤40nm by an oligonucleotide mediated ligation and enzymatic amplification reaction whose product is subsequently recognized by a fluorescent probe. Consequently, each fluorescent spot indicates that two proteins are in proximity. The mouse/rabbit Duolink in situ red starter kit (Olink, Uppsala, Sweden) was used according to the manufacturer’s instructions. MDA-MB-231 cells were seeded at 1×10^4^ cells per well in a 6-well plate. The cells were fixed and permeabilized as described above for immunofluorescence studies. Then, the cells were incubated with a 1:100 dilution of the primary antibodies (RhoA and DIA) in PBS containing 0.01% BSA overnight at 4°C. For the rest of the protocol the manufacturer’s instructions were followed. Briefly, the cells were washed in Buffer A (supplied with the kit) 3 times for 15 minutes and incubated with the PLA probes for one hour at 37°C in a humidified chamber. This was followed by a 10 minute and a 5 minute wash in Buffer A. The ligation reaction was carried out at 37°C for one hour in a humidified chamber followed by a 10 and 5 minute wash in Buffer A. The cells were then incubated with the amplification mix for two hours at 37°C in a darkened humidified chamber. After washing with 1x Buffer B (supplied with the kit) for 10 minutes and 1 minute wash with 0.01x buffer B, followed by 488 phalloidin staining (Molecular Probes Catalog number: A12379) to visualize cellular F-actin, the cells were mounted using the mounting media (containing DAPI to visualize cell nucleus) supplied with the kit. Images were quantified using Fiji distribution of ImageJ. A longitudinal axis emanating at the cell front was drawn through selected cells. Along this axis the cell was divided into 3 segments: 10% corresponding to the cell front, 70% corresponding to the cell middle, and 20% corresponding to the cell rear. Then the image was converted into a 2-bit image and masks over PLA reactions were drawn. Finally, the number of PLA reactions per segment as well as the total area occupied by PLA signals per segment were quantified. All the statistical analysis for PLA was done in Excel.

##### Random migration assays

Cell migration assays were performed with cell lines stably expressing nuclear mKATE2 (a red fluorescence protein allowing cell tracking) treated with either vehicle, the ROCK inhibitors Y-27632 (10μM) or GSK-269962 (1μM). Cells were seeded on IncuCyte® ImageLock 96-well plates (cat.4379) at 100 cells per well and placed into an IncuCyte™ ZOOM with a dual color filter unit. Images were captured every 10 min using phase contrast and red channel with an 10×/0.25 ph1 objective, over a 24 h period. Stacks of the red florescence channel were created. ImageJ software was used to enhance contrast, subtract background and transform the images to 8-bit greyscale. Random migration trajectories were obtained from the images using the FastTracks Matlab plugin ^66^, subsequent statistical analysis and plotting were done in Python.

##### Assaying RhoA activity by live-cell FRET imaging

MDA-MB-231 stably expressing the mTFP-YFP RhoA-GTP FRET biosensor ^41^ were seeded in Fluorodish™ glass-bottomed plate (cat.FD35-100) coated with collagen. Cells were treated as indicated for siRNA or ROCK inhibition experiments (Y-27632 2.5μM). The biosensor-expressing cells were imaged at 5-sec intervals for 10min in an Andor Dragonfly spinning disk confocal microscope with a 60x/1.4 - Oil objective. An excitation wavelength (445 nm) was used for both mTFP and FRET channels, while 480 and 540 nm emission filters were used for the mTFP and FRET channels, respectively, with the Confocal 40μm High Sensitivity imaging mode. A cell-free area using the same settings for exposure and time was acquired for background correction. The raw images were de-noised with the ImageJ PureDenoise plugin ^67^, and ratiometric images were generated. Kymographs were built using MultiKymographr plugin.

##### Modeling

###### Numerical methods for solving PDE equations

The PDE system (Eq. 10) was solved numerically by the finite volume method ^68^ aided by the splitting technique ^69^, and using the OpenFOAM platform ^70^. A computational 2D domain was obtained by extracting contours of cells from experimental cell images using the OpenCV library ^71^ and meshed by non-structured triangular meshes using the Salome platform ^72^. An example of the computational mesh is presented in Fig. S4F. The x and y axes were set along the cell length and width as depicted in Fig. 3A. Distributions of the total concentrations of DIA and ROCK were set according to Eqs. 3 and 10. For equations describing spatiotemporal dynamics of active and inactive forms of Rho and Rac1, zero-gradient boundary conditions were applied. The diffusion term was discretized using unstructured triangular meshes by means of the “over-relaxed correction” technique ^73^. ODE systems describing chemical kinetics were solved using fifth-order Cash-Karp embedded Runge-Kutta scheme with error control and adjusted time-step size ^74^. The simulation results were visualized using the ParaView software package ^75^.

## Supporting information

Supplemental Text, Tables, Figiure and Movie Legends

## Acknowledgments

We thank Dirk Fey (Systems Biology Ireland, University College Dublin) for his input in the modeling and discussions, Kasia Kedziora (Netherlands Cancer Institute) for providing macros for quantifying the PLA images, and Denis Pushin (Moscow Institute of Physics and Technology) for advices on obtaining the contours of cells from the experimental images for carrying out numerical calculations. We thank the Edinburgh Super-Resolution Imaging Consortium for assistance with super-resolution imaging. Supported by NIH/NCI grant R01CA244660, EU grants SmartNanoTox (grant no. 686098), NanoCommons (grant no. 731032), SFI grants 14/IA/2395 and 18/SPP/3522, CRUK Edinburgh Centre C157/A25140, Breast Cancer NOW PR183.

## Author Contributions

B.N.K. conceived the study. O.S.R., M.A.T, E.N. and B.N.K. developed the model with input from W.K. AvK designed the experiments with input from W.K. and O.S.R. A.B-C., A.W., E.N., A.M.G. and AvK conducted the experiments. B.N.K., O.S.R. and W.K. wrote the manuscript with input from all authors.

