## Supplemental Text, Tables, Figiure and Movie Legends for "Periodic propagating waves coordinate RhoGTPase network dynamics at the leading and trailing edges during cell migration"

**Supplemental Methods**

**Detailed description of the mathematical model**

**Section 1. Relating the PLA data to the total effector concentrations.**

The PLA data showed that RhoA interactions with its effectors DIA and ROCK change along the cell from the cell rear to the leading edge (Figs. 1A and 1B). This correlates with our experimental data (Figs. 1C and 1D) and the literature data on DIA and ROCK localization, suggesting that the concentrations of DIA and ROCK are different at the leading edge, in the middle of the cell, and at the cell rear ^1-5^. The steady-state concentration of the complex of RhoA-GTP $\left( \left[ Rho\text{-}T \right] \right)$ and DIA $\left( \left[ DIA\text{-}Rho\text{-}T \right] \right)$ can be derived using the rapid equilibrium approximation and the dissociation constant ($K_{d}^{RhoDIA}$). Taking into account the moiety conservation for DIA, we obtain,

$$\left[ DIA \right]\cdot\left[ Rho\text{-}T \right]=K_{d}^{RhoDIA}\cdot\left[ DIA\text{-}Rho\text{-}T \right]$$

$${DIA}^{tot}=\left[ DIA \right]+\left[ DIA\text{-}Rho\text{-}T \right]$$

(1).

Our quantitative proteomic data suggest that the RhoA abundance is at least 10-fold higher than the abundance of all DIA isoforms combined, Table S1 ^6^. Therefore, in Eq. 1 we can neglect the changes in the RhoA-GTP concentration caused by the RhoA-GTP sequestration into the complex with DIA. The $K_{d}^{RhoDIA}$ is at least two orders of magnitude smaller than the RhoA abundance ^7^, which leads to an approximate, linear dependence of the complex concentration on the total DIA abundance

$$\left[ DIA\text{-}Rho\text{-}T \right]=\frac{DIA^{tot}\cdot\left[ Rho\text{-}T \right]}{K_{d}^{RhoDIA}+\left[ Rho\text{-}T \right]} \sim DIA^{tot}$$

(2).

Thus, our data on the changes in the RhoA-DIA complexes along the cell length at the constant RhoA-GTP level can be interpreted as the changes in the abundance of DIA that can bind RhoA-GTP in the plasma membrane, corroborating the literature data ^1-5^.

The abundance of all ROCK isoforms is also much smaller than the RhoA abundance (see Table S1), which together with the cooperative binding of ROCK domains to active RhoA ^8^ allows us to conclude that the RhoA-GTP-ROCK complex concentration can also be approximated as a linear function of the total ROCK abundance (${ROCK}^{tot}$). Consequently, in the model the total abundances of DIA and ROCK depend on the spatial coordinate along the cell, as shown in Figs. 3B and 3C. Associating the x axis with the cell length and considering the y axis along the cell width, we use the following distribution of the DIA and ROCK abundances along the x-axis,

$$DIA^{tot}\left( x \right)=\left( DIA_{h}-DIA_{l} \right)\cdot\frac{x}{L}+DIA_{l}, DIA_{h}>DIA_{l}$$

$${ROCK}^{tot}\left( x \right)=\left\{ \begin{aligned} ROCK_{l}, 0\leq x\leq x_{l} \\ ROCK_{h}, x_{l}\leq x\leq L \end{aligned} \right., ROCK_{h}>ROCK_{l}$$

(3).

where $L$ is the cell length.

**Section 2. Modeling the RhoA - Rac1 network dynamics**

The spatiotemporal dynamics of the RhoA - Rac1 network are governed by a partial differential equation (PDE) system, referred to as a reaction-diffusion model. To derive this PDE system, we first consider ordinary differential equation (ODE) systems that describe biochemical reactions and RhoA and Rac1 interactions with their effectors at any fixed point in the cellular space. The difference between the ODE systems at distinct spatial points is brought about by the changes in the total abundances of ROCK1 and DIA along the longitudinal axis of polarized cells given by Eq. 3 (see also Figs. 3B and 3C). These ODE equations are then converted to a PDE system by accounting for the diffusion fluxes of active and inactive protein forms.

The model was populated by the protein abundances from our quantitative mass spectrometry data ^6^. The data suggested that Rac1 and RhoA were the most abundant Rac and Rho isoforms and that their levels exceed the abundances of PAK, ROCK and DIA isoforms combined by an order of magnitude (Table S1). The abundances of ROCK1 and ROCK2 were comparable, DIA1 was the most abundant DIA isoform, and PAK2 was the only detected PAK isoform.

We considered the time scale on which the total abundances of RhoA (${Rho}^{tot}$), DIA (${DIA}^{tot}$), ROCK (${ROCK}^{tot}$), Rac1 (${Rac}^{tot}$) and PAK (${PAK}^{tot}$) are conserved. We denote active, GTP-bound forms of RhoA and Rac1 by $\left[ Rho\text{-}T \right]$ and $\left[ Rac\text{-}T \right]$, and inactive GDP-bound forms by $\left[ Rho\text{-}D \right]$ and $\left[ Rac\text{-}D \right]$. Active forms of DIA, ROCK and active (phosphorylated) PAK are denoted by $\left[ DIA^{*} \right]$, $\left[ ROCK^{*} \right]$ and $\left[ pPAK \right],$ respectively. Because of the conservation constraints, the concentrations of active forms can be approximately expressed as the corresponding total abundances minus concentrations of inactive forms. Then, assuming the Michaelis-Menten kinetics for the rates of activation and deactivation reactions of the active forms of the GTPases and their effectors ($\left[ DIA^{*} \right]$, $\left[ ROCK^{*} \right]$ and $\left[ pPAK \right]$), the temporal kinetics of the network are given by the following system of ODEs,

$\frac{d\left[ Rho\text{-}T \right]}{dt}=\alpha_{DIA}^{Rho}\alpha_{PAK}^{Rho}V_{GEF}^{Rho}\frac{\left( Rho^{tot}-\left[ Rho\text{-}T \right] \right)/{K_{GEF}^{Rho}}}{1+\left( Rho^{tot}-\left[ Rho\text{-}T \right] \right)/{K_{GEF}^{Rho}}}-V_{GAP}^{Rho}\frac{\left[ Rho\text{-}T \right]/{K_{GAP}^{Rho}}}{1+\left[ Rho\text{-}T \right]/{K_{GAP}^{Rho}}}$

$\frac{d\left[ DIA^{*} \right]}{dt}=\alpha_{Rho}^{DIA}V_{a}^{DIA}\frac{\left( DIA^{tot}-\left[ DIA^{*} \right] \right)/{K_{a}^{DIA}}}{1+\left( DIA^{tot}-\left[ DIA^{*} \right] \right)/{K_{a}^{DIA}}}-V_{i}^{DIA}\frac{\left[ DIA^{*} \right]/{K_{i}^{DIA}}}{1+\left[ DIA^{*} \right]/{K_{i}^{DIA}}}$

$\frac{d\left[ {ROCK}^{*} \right]}{dt}=\alpha_{Rho}^{ROCK}V_{a}^{ROCK}\frac{\left( {ROCK}^{tot}-\left[ {ROCK}^{*} \right] \right)/{K_{a}^{ROCK}}}{1+\left( {ROCK}^{tot}-\left[ {ROCK}^{*} \right] \right)/{K_{a}^{ROCK}}}-V_{i}^{ROCK}\frac{\left[ {ROCK}^{*} \right]/{K_{i}^{ROCK}}}{1+\left[ {ROCK}^{*} \right]/{K_{i}^{ROCK}}}$

$\frac{d\left[ Rac\text{-}T \right]}{dt}=\alpha_{DIA}^{Rac}\alpha_{PAK}^{Rac}V_{GEF}^{Rac}\frac{\left( {Rac}^{tot}-\left[ Rac\text{-}T \right] \right)/{K_{GEF}^{Rac}}}{1+\left( {Rac}^{tot}-\left[ Rac\text{-}T \right] \right)/{K_{GEF}^{Rac}}}-\alpha_{ROCK}^{Rac}V_{GAP}^{Rac}\frac{\left[ Rac\text{-}T \right]/{K_{GAP}^{Rac}}}{1+\left[ Rac\text{-}T \right]/{K_{GAP}^{Rac}}}$

$\frac{d\left[ pPAK \right]}{dt}=\alpha_{Rac}^{PAK}V_{a}^{PAK}\frac{\left( {PAK}^{tot}-\left[ pPAK \right] \right)/{K_{a}^{PAK}}}{1+\left( {PAK}^{tot}-\left[ pPAK \right] \right)/{K_{a}^{PAK}}}-V_{i}^{PAK}\frac{\left[ pPAK \right]/{K_{i}^{PAK}}}{1+\left[ pPAK \right]/{K_{i}^{PAK}}}$

(4).

Here the maximal rates and the Michaelis-Menten constants are denoted by the capital letters V’s and K’s with relevant indices. These V’s values correspond to the maximal rates in the absence of positive or negative regulatory interactions between GTPases, which modify reaction rates. We describe the regulatory interactions, which specify the negative or positive influence of the active form of protein *Y* on protein *X*, by the dimensionless multipliers $\alpha_{Y}^{X}$ (illustrated in Fig. S2E) ^9^. Assuming general hyperbolic modifier kinetics, each multiplier $\alpha_{Y}^{X}$ has the same functional form ^10^,

$$\alpha_{Y}^{X}=\frac{1+\gamma_{Y}^{X}\cdot Y_{a}/K_{Y}^{X}}{1+Y_{a}/K_{Y}^{X}}$$

(5).

Here $Y_{a}$ is active form of protein *Y*. The coefficient $\gamma_{Y}^{X}$ > 1 indicates activation; $\gamma_{Y}^{X}$< 1 inhibition; and $\gamma_{Y}^{X}$ = 1 denotes the absence of regulatory interactions, in which case the modifying multiplier $\alpha_{Y}^{X}$ equals 1. $K_{Y}^{X}$ is the activation or inhibition constant.

***Model-predicted different temporal dynamics of the GTPase activities***

Substituting the expressions for modifying multipliers (Eq. 5) into Eqs. 4, we obtain the following equations governing the temporal dynamics of the active protein forms.

$\frac{d\left[ Rho\text{-}T \right]}{dt}=V_{GEF}^{Rho}\frac{1+{\gamma_{DIA}^{Rho}\left[ DIA^{*} \right]}/{K_{DIA}^{Rho}}}{1+\left[ DIA^{*} \right]/{K_{DIA}^{Rho}}}\frac{1+{\gamma_{PAK}^{Rho}\left[ pPAK \right]}/{K_{PAK}^{Rho}}}{1+\left[ pPAK \right]/{K_{PAK}^{Rho}}}\frac{\left( Rho^{tot}-\left[ Rho\text{-}T \right] \right)/{K_{GEF}^{Rho}}}{1+\left( Rho^{tot}-\left[ Rho\text{-}T \right] \right)/{K_{GEF}^{Rho}}}$

$-V_{GAP}^{Rho}\frac{\left[ Rho\text{-}T \right]/{K_{GAP}^{Rho}}}{1+\left[ Rho\text{-}T \right]/{K_{GAP}^{Rho}}}$

$\frac{d\left[ DIA^{*} \right]}{dt}=V_{a}^{DIA}\frac{1+{\gamma_{Rho}^{DIA}\left[ Rho\text{-}T \right]}/{K_{Rho}^{DIA}}}{1+\left[ Rho\text{-}T \right]/{K_{Rho}^{DIA}}}\frac{\left( DIA^{tot}-\left[ DIA^{*} \right] \right)/{K_{a}^{DIA}}}{1+\left( DIA^{tot}-\left[ DIA^{*} \right] \right)/{K_{a}^{DIA}}}-V_{i}^{DIA}\frac{\left[ DIA^{*} \right]/{K_{i}^{DIA}}}{1+\left[ DIA^{*} \right]/{K_{i}^{DIA}}}$

$\frac{d\left[ {ROCK}^{*} \right]}{dt}=V_{a}^{ROCK}\frac{1+{\gamma_{Rho}^{ROCK}\left[ Rho\text{-}T \right]}/{K_{Rho}^{ROCK}}}{1+\left[ Rho\text{-}T \right]/{K_{Rho}^{ROCK}}}\frac{\left( {ROCK}^{tot}-\left[ {ROCK}^{*} \right] \right)/{K_{a}^{ROCK}}}{1+\left( {ROCK}^{tot}-\left[ {ROCK}^{*} \right] \right)/{K_{a}^{ROCK}}}$

$-V_{i}^{ROCK}\frac{\left[ {ROCK}^{*} \right]/{K_{i}^{ROCK}}}{1+\left[ {ROCK}^{*} \right]/{K_{i}^{ROCK}}}$

$\frac{d\left[ Rac\text{-}T \right]}{dt}=V_{GEF}^{Rac}\frac{1+{\gamma_{DIA}^{Rac}\left[ DIA^{*} \right]}/{K_{DIA}^{Rac}}}{1+\left[ DIA^{*} \right]/{K_{DIA}^{Rac}}}\frac{1+{\gamma_{PAK}^{Rac}\left[ pPAK \right]}/{K_{PAK}^{Rac}}}{1+\left[ pPAK \right]/{K_{PAK}^{Rac}}}\frac{\left( {Rac}^{tot}-\left[ Rac\text{-}T \right] \right)/{K_{GEF}^{Rac}}}{1+\left( {Rac}^{tot}-\left[ Rac\text{-}T \right] \right)/{K_{GEF}^{Rac}}}$

$-V_{GAP}^{Rac}\frac{1+{\gamma_{ROCK}^{Rac}\left[ {ROCK}^{*} \right]}/{K_{ROCK}^{Rac}}}{1+\left[ {ROCK}^{*} \right]/{K_{ROCK}^{Rac}}}\frac{\left[ Rac\text{-}T \right]/{K_{GAP}^{Rac}}}{1+\left[ Rac\text{-}T \right]/{K_{GAP}^{Rac}}}$

$\frac{d\left[ pPAK \right]}{dt}=V_{a}^{PAK}\frac{1+{\gamma_{Rac}^{PAK}\left[ Rac\text{-}T \right]}/{K_{Rac}^{PAK}}}{1+\left[ Rac\text{-}T \right]/{K_{Rac}^{PAK}}}\frac{\left( {PAK}^{tot}-\left[ pPAK \right] \right)/{K_{a}^{PAK}}}{1+\left( {PAK}^{tot}-\left[ pPAK \right] \right)/{K_{a}^{PAK}}}$

$-V_{i}^{PAK}\frac{\left[ pPAK \right]/{K_{i}^{PAK}}}{1+\left[ pPAK \right]/{K_{i}^{PAK}}}$

(6).

Because $DIA^{tot}$ and ${ROCK}^{tot}$ depend on the spatial coordinate along the cell (Eq. 3), and $DIA^{tot}$, ${ROCK}^{tot}$, and ${PAK}^{tot}$ were perturbed experimentally, we first explored the different possible types of the network temporal dynamics (Eq. 6) in the parameter space of these three effector abundances. We obtained bifurcation diagrams in each of the three planes of the two effector abundances and classified different types of the dynamic regimes that can be detected (Figs. 2C, 4A, S2A-C). We used BioNetGen ^11^^,^^12^ and DYVIPAC ^13^, software packages, and SciPy ^14^ and Matplotlib Python libraries ^15^. The code that performs calculations and plotting can be found in the Supplemental Information.

To get initial insights into different dynamic regimes of this five ODE system (Eq. 6), we analyzed the vector fields and the nullclines for a two ODE system, obtained using the quasi steady-state approximation. Because the concentrations of active forms of DIA, ROCK and PAK are an order of magnitude less than the GTPase concentrations, we can express these active effector concentrations in terms of $\left[ Rho\text{-}T \right]$ and $\left[ Rac\text{-}T \right]$ by applying the quasi steady-state approximation, as follows ^9^,

$\left\{ \begin{aligned} \frac{d\left[ DIA^{*} \right]}{dt}=0 \\ \frac{d\left[ {ROCK}^{*} \right]}{dt}=0 \\ \frac{d\left[ pPAK \right]}{dt}=0 \end{aligned} \right. \to$ $\left\{ \begin{aligned} \left[ DIA^{*} \right]=f_{DIA}\left( \left[ Rho\text{-}T \right], \left[ Rac\text{-}T \right] \right) \\ \left[ ROCK^{*} \right]=f_{ROCK}\left( \left[ Rho\text{-}T \right], \left[ Rac\text{-}T \right] \right) \\ \left[ pPAK \right]=f_{PAK}\left( \left[ Rho\text{-}T \right], \left[ Rac\text{-}T \right] \right) \end{aligned} \right.$

(7).

To find the functions, $f_{DIA}$, $f_{ROCK}$, and $f_{PAK}$, Eq. 7 were solved numerically for each value of active RhoA and Rac1. The solutions were substituted into the equations governing the dynamics of RhoA-GTP and Rac1-GTP (see Eq. 6) to obtain the following system of only two differential equations.

$\frac{d\left[ Rho\text{-}T \right]}{dt}=V_{GEF}^{Rho}\frac{1+{\gamma_{DIA}^{Rho}f_{DIA}}/{K_{DIA}^{Rho}}}{1+{f_{DIA}}/{K_{DIA}^{Rho}}}\frac{1+{\gamma_{PAK}^{Rho}f_{PAK}}/{K_{PAK}^{Rho}}}{1+{f_{PAK}}/{K_{PAK}^{Rho}}}\frac{\left( Rho^{tot}-\left[ Rho\text{-}T \right] \right)/{K_{GEF}^{Rho}}}{1+\left( Rho^{tot}-\left[ Rho\text{-}T \right] \right)/{K_{GEF}^{Rho}}}$

$-V_{GAP}^{Rho}\frac{\left[ Rho\text{-}T \right]/{K_{GAP}^{Rho}}}{1+\left[ Rho\text{-}T \right]/{K_{GAP}^{Rho}}}$

$\frac{d\left[ Rac\text{-}T \right]}{dt}=V_{GEF}^{Rac}\frac{1+{\gamma_{DIA}^{Rac}f_{DIA}}/{K_{DIA}^{Rac}}}{1+{f_{DIA}}/{K_{DIA}^{Rac}}}\frac{1+{\gamma_{PAK}^{Rac}f_{PAK}}/{K_{PAK}^{Rac}}}{1+{f_{PAK}}/{K_{PAK}^{Rac}}}\frac{\left( {Rac}^{tot}-\left[ Rac\text{-}T \right] \right)/{K_{GEF}^{Rac}}}{1+\left( {Rac}^{tot}-\left[ Rac\text{-}T \right] \right)/{K_{GEF}^{Rac}}}$

$-V_{GAP}^{Rac}\frac{1+{\gamma_{ROCK}^{Rac}f_{ROCK}}/{K_{ROCK}^{Rac}}}{1+{f_{ROCK}}/{K_{ROCK}^{Rac}}}\frac{\left[ Rac\text{-}T \right]/{K_{GAP}^{Rac}}}{1+\left[ Rac\text{-}T \right]/{K_{GAP}^{Rac}}}$

(8).

Figures S3A-S3I illustrate the vector fields and nullclines for a 2-D system describing the temporal dynamics of RhoA-GTP and Rac1-GTP. Each dynamic regime shown in Figs. 2C, 4A and S2A-S2C has the corresponding phase portrait in Fig. S3. The red line represents the solution for the equation $d\left[ Rho\text{-}T \right]/dt=0$ (the RhoA nullcline), and the blue line represents the solution for the equation $d\left[ Rac\text{-}T \right]/dt=0$ (the Rac1 nullcline).

Points of intersection of the nullclines are network steady states for both 5 ODE and 2 ODE systems. These states can be stable or unstable (shown by bold points or triangles, respectively in Figs. S3A-S3I). For each of dynamic regimes 0, 1 and 6 there is only a single steady state, which is a stable focus for regime 0, stable node for regime 6 and an unstable focus for regime 1 (points 1 at Figs. S3A, S3B and S3G). If a steady state is unstable focus, self-sustained oscillations (a limit cycle) may or may not exist in the system, depending on the global topology of the vector fields. In our system, although unstable focus steady states are observed in regimes 1 - 5 and 7, self-sustained oscillations exist only in regimes 1 and 3. For these oscillatory regimes, we plotted projections of the limit cycle trajectory calculated for a 5-dimensional ODE system (Eq. 6) to a 2-dimensional space of active RhoA and active Rac1 concentrations (green curves in Figs. S3B and S3D).

The increase in the DIA abundance at low, fixed ROCK abundance can transform dynamic regime 0 into dynamic regime 1 (Fig. S2A) following the Andronov-Hopf bifurcation ^16^. This bifurcation results in losing the stability of the focus (point 1, Figs. S3A and S3B) and the appearance of a limit cycle around the unstable focus (green trajectory, Fig. S3B).

The increase in ROCK abundance at fixed DIA abundance can transform dynamic regime 1 into regime 3 termed BiDR (Fig. 2C). At certain increased ROCK abundances, the Rac1 nullcline crosses the RhoA nullcline generating a saddle point and a stable node (points 2 and 3, Fig S3D), known as a saddle-node bifurcation ^16^. In the BiDR regime a stable limit cycle coexists with a stable node, and each of these dynamic regimes has its own basin of attraction (Fig. S3D). A saddle point separates the basins of attraction of the limit cycle and the stable node. The further increase in the ROCK abundance moves the system to regime 2 where the limit cycle disappears, whereas an unstable focus (point 1, Fig. S3C), saddle (point 2, Fig. S3C) and stable node (point 3, Fig. S3C) persist. The disappearance of the limit cycle occurs when it merges with a saddle point in the process termed as a saddle homoclinic bifurcation ^17^. Thus, although regimes 2 and 3 have the same number and stability types of the steady-state solutions, a stable limit cycle exists only in regime 3.

If the DIA abundance increases at the high, fixed ROCK abundance, a saddle-node bifurcation appears earlier than the Andronov-Hopf bifurcation, and dynamic regime 1 with single stable focus transforms into dynamic regime 4 (Fig. S2A) with stable node (point 3, Fig. S3E) and saddle point (point 2, Fig. S3E) in addition to the stable focus (point 1, Fig. S3E). At the point where dynamic regimes 0 - 4 converge, the saddle-node, the saddle homoclinic and the Andronov-Hopf bifurcations happen simultaneously in a process known as the Bogdanov-Takens bifurcation ^16^.

Regimes 4 and 8 have two stable steady states (points 1 and 3, Figs. S3E and S3I) and one saddle point (point 2, Figs. S3E and S3I), which separates the basins of attraction of the stable states. Regime 8 is a classic bistability regime arising from a double negative feedback in the RhoA-Rac1 network. One stable node has the high RhoA and low Rac activities, whereas the other stable node has the high Rac and low Rho activities (points 1 and 3, Fig. S3I) ^6^. In regime 4 one of the stable steady states is a stable node, whereas the other is a stable focus. Both stable states have low Rac1-GTP levels, but the stable focus (point 1, Fig. S3E) has a low RhoA-GTP level, while the stable node (point 3, Fig. S3E) has a high RhoA-GTP level. Regime 4 occurs for low DIA abundances, when the activating connection from RhoA to Rac1 is weak. The dynamical behavior of regime 7 is similar to the dynamics of regime 8. Both regimes exhibit two stable nodes (points 3 and 5 for regime 7, Fig. S3H) and a saddle resulting in bistability. Regime 7 has an additional unstable focus and saddle (points 1-2, Fig. S3H), which do not substantially change the basins of attraction of stable nodes.

Regime 6 has a single steady state that is a stable node, to which all solutions converge regardless of the initial conditions (Fig. S3G). The dynamical behavior of regime 5 is similar to the dynamics of regime 6. Regime 5 has a single stable node but also an additional unstable focus and saddle (points 1-2, Fig. S3F), which does not substantially change the basin of attraction of the stable node.

Summarizing, the above analysis of a 2-D system (Eq. 8) helped us comprehend the dynamic behaviors and parameter bifurcation diagrams obtained for a 5-D system (Eq. 6, in Figs. 2C, 4A and S2A-S2C).

***Describing spatiotemporal dynamical regimes in the model***

To explore the spatiotemporal behavior of the RhoA-Rac1 network in an entire cell we took into account diffusion fluxes and spatial distribution of RhoA, Rac1 and their effectors. The spatiotemporal dynamics of the system is described by the following system of partial differential equations (PDEs). Since active and inactive forms of RhoA and Rac1 GTPases can have different diffusion coefficients, the PDEs include both protein forms.

$\frac{\partial\left[ Rho\text{-}T \right]}{\partial t}=V_{GEF}^{Rho}\frac{1+{\gamma_{DIA}^{Rho}\left[ DIA^{*} \right]}/{K_{DIA}^{Rho}}}{1+\left[ DIA^{*} \right]/{K_{DIA}^{Rho}}}\frac{1+{\gamma_{PAK}^{Rho}\left[ pPAK \right]}/{K_{PAK}^{Rho}}}{1+\left[ pPAK \right]/{K_{PAK}^{Rho}}}\frac{\left[ Rho\text{-}D \right]/{K_{GEF}^{Rho}}}{1+\left[ Rho\text{-}D \right]/{K_{GEF}^{Rho}}}$

$-V_{GAP}^{Rho}\frac{\left[ Rho\text{-}T \right]/{K_{GAP}^{Rho}}}{1+\left[ Rho\text{-}T \right]/{K_{GAP}^{Rho}}}-\nabla\left( -D_{RhoT}\nabla\left[ Rho\text{-}T \right] \right)$

$\frac{\partial\left[ Rho\text{-}D \right]}{\partial t}=-V_{GEF}^{Rho}\frac{1+{\gamma_{DIA}^{Rho}\left[ DIA^{*} \right]}/{K_{DIA}^{Rho}}}{1+\left[ DIA^{*} \right]/{K_{DIA}^{Rho}}}\frac{1+{\gamma_{PAK}^{Rho}\left[ pPAK \right]}/{K_{PAK}^{Rho}}}{1+\left[ pPAK \right]/{K_{PAK}^{Rho}}}\frac{\left[ Rho\text{-}D \right]/{K_{GEF}^{Rho}}}{1+\left[ Rho\text{-}D \right]/{K_{GEF}^{Rho}}}$

$+V_{GAP}^{Rho}\frac{\left[ Rho\text{-}T \right]/{K_{GAP}^{Rho}}}{1+\left[ Rho\text{-}T \right]/{K_{GAP}^{Rho}}}-\nabla\left( -D_{RhoD}\nabla\left[ Rho\text{-}D \right] \right)$

$\frac{\partial\left[ DIA^{*} \right]}{\partial t}=V_{a}^{DIA}\frac{1+{\gamma_{Rho}^{DIA}\left[ Rho\text{-}T \right]}/{K_{Rho}^{DIA}}}{1+\left[ Rho\text{-}T \right]/{K_{Rho}^{DIA}}}\frac{\left( DIA^{tot}\left( \vec{x} \right)-\left[ DIA^{*} \right] \right)/{K_{a}^{DIA}}}{1+\left( DIA^{tot}\left( \vec{x} \right)-\left[ DIA^{*} \right] \right)/{K_{a}^{DIA}}}-V_{i}^{DIA}\frac{\left[ DIA^{*} \right]/{K_{i}^{DIA}}}{1+\left[ DIA^{*} \right]/{K_{i}^{DIA}}}$

$\frac{\partial\left[ {ROCK}^{*} \right]}{\partial t}=V_{a}^{ROCK}\frac{1+{\gamma_{Rho}^{ROCK}\left[ Rho\text{-}T \right]}/{K_{Rho}^{ROCK}}}{1+\left[ Rho\text{-}T \right]/{K_{Rho}^{ROCK}}}\frac{\left( {ROCK}^{tot}\left( \vec{x} \right)-\left[ {ROCK}^{*} \right] \right)/{K_{a}^{ROCK}}}{1+\left( {ROCK}^{tot}\left( \vec{x} \right)-\left[ {ROCK}^{*} \right] \right)/{K_{a}^{ROCK}}}$

$-V_{i}^{ROCK}\frac{\left[ {ROCK}^{*} \right]/{K_{i}^{ROCK}}}{1+\left[ {ROCK}^{*} \right]/{K_{i}^{ROCK}}}$

$\frac{\partial\left[ Rac\text{-}T \right]}{\partial t}=V_{GEF}^{Rac}\frac{1+{\gamma_{DIA}^{Rac}\left[ DIA^{*} \right]}/{K_{DIA}^{Rac}}}{1+\left[ DIA^{*} \right]/{K_{DIA}^{Rac}}}\frac{1+{\gamma_{PAK}^{Rac}\left[ pPAK \right]}/{K_{PAK}^{Rac}}}{1+\left[ pPAK \right]/{K_{PAK}^{Rac}}}\frac{\left[ Rac\text{-}D \right]/{K_{GEF}^{Rac}}}{1+\left[ Rac\text{-}D \right]/{K_{GEF}^{Rac}}}$

$-V_{GAP}^{Rac}\frac{1+{\gamma_{ROCK}^{Rac}\left[ {ROCK}^{*} \right]}/{K_{ROCK}^{Rac}}}{1+\left[ {ROCK}^{*} \right]/{K_{ROCK}^{Rac}}}\frac{\left[ Rac\text{-}T \right]/{K_{GAP}^{Rac}}}{1+\left[ Rac\text{-}T \right]/{K_{GAP}^{Rac}}}-\nabla\left( -D_{RacT}\nabla\left[ Rac\text{-}T \right] \right)$

$\frac{\partial\left[ Rac\text{-}D \right]}{\partial t}=-V_{GEF}^{Rac}\frac{1+{\gamma_{DIA}^{Rac}\left[ DIA^{*} \right]}/{K_{DIA}^{Rac}}}{1+\left[ DIA^{*} \right]/{K_{DIA}^{Rac}}}\frac{1+{\gamma_{PAK}^{Rac}\left[ pPAK \right]}/{K_{PAK}^{Rac}}}{1+\left[ pPAK \right]/{K_{PAK}^{Rac}}}\frac{\left[ Rac\text{-}D \right]/{K_{GEF}^{Rac}}}{1+\left[ Rac\text{-}D \right]/{K_{GEF}^{Rac}}}$

$+V_{GAP}^{Rac}\frac{1+{\gamma_{ROCK}^{Rac}\left[ {ROCK}^{*} \right]}/{K_{ROCK}^{Rac}}}{1+\left[ {ROCK}^{*} \right]/{K_{ROCK}^{Rac}}}\frac{\left[ Rac\text{-}T \right]/{K_{GAP}^{Rac}}}{1+\left[ Rac\text{-}T \right]/{K_{GAP}^{Rac}}}-\nabla\left( -D_{RacD}\nabla\left[ Rac\text{-}D \right] \right)$

$\frac{\partial\left[ pPAK \right]}{\partial t}=V_{a}^{PAK}\frac{1+{\gamma_{Rac}^{PAK}\left[ Rac\text{-}T \right]}/{K_{Rac}^{PAK}}}{1+\left[ Rac\text{-}T \right]/{K_{Rac}^{PAK}}}\frac{\left( {PAK}^{tot}-\left[ pPAK \right] \right)/{K_{a}^{PAK}}}{1+\left( {PAK}^{tot}-\left[ pPAK \right] \right)/{K_{a}^{PAK}}}$

$-V_{i}^{PAK}\frac{\left[ pPAK \right]/{K_{i}^{PAK}}}{1+\left[ pPAK \right]/{K_{i}^{PAK}}}$

(9).

Here, $D_{RhoT}$ and $D_{RhoD}$ are the diffusion coefficients of active and inactive forms of RhoA, and $D_{RacT}$ and $D_{RacD}$ are the diffusion coefficients of active and inactive forms of Rac1. For all forms of RhoA and Rac1, zero-gradient boundary conditions are considered at the boundaries of the computational domain, describing no flux conditions at the cell borders. The spatial profiles of the total DIA and ROCK concentrations are set by Eq. 3.

At the leading edge, the total concentrations of DIA and ROCK correspond to oscillatory regimes 1 and 3 observed for a well-mixed system (Figs. 2A, 2B, 2F, S3B and S3D). In the spatial case, the PDE equations (Eq. 9) with these parameters generate excitable media, where self-sustained waves of the RhoA and Rac1 activities are formed periodically. Thus, the leading edge can be considered as a “pacemaker” of the GTPase cellular machinery ^18^, by analogy to the sinoatrial node in the heart ^19^.

At the cell body and rear the total concentration of DIA is lower, and the total concentration of ROCK is higher than at the leading edge. For the well-mixed system (Eq. 6), these concentration parameters correspond to regime 2 (Fig. 2A, 2C and S3C). For the dynamics in space and time, these parameters bring about weakly excitable media, which can propagate self-sustained waves of RhoA and Rac1 activities after receiving a strong stimulus, but unable to autonomously generate such waves. In the stimulus absence, high RhoA and low Rac1 stationary activities are maintained in this media. Following an over-threshold stimulus, this weakly excitable media propagates the wave of high Rac1 activity, and then returns to the steady state with high RhoA and low Rac1 activities. Importantly, the excitability of this media gradually decays approaching the cell rear. As a result, in a mesenchymal polarized cell a number of waves of RhoA and Rac1 activity must be generated at the leading edge to induce a self-sustained wave in the cell body and rear, in contrast with the heart where every wave generated in the sinoatrial node spreads through the entire heart. The higher concentration of ROCK exists at the cell body and rear, the higher number of waves must be generated at the leading edge before a GTPase activity wave propagates through an entire cell. If the total ROCK concentration of is too high in the cell body and rear, the waves generated at the leading edge vanish before propagating deeply into the cell and reaching the cell rear.

Thus, high excitability at the leading edge and low excitability in the cell body and at the rear result in a cyclic dynamic pattern, in which multiple protrusion-retraction cycles are generated at the leading edge before a migrating cell moves.

***Modeling the mechanisms of PAK and ROCK inhibition***

The mechanism of PAK inhibition by allosteric inhibitor IPA-3 was modeled similarly as in our previous study ^6^. IPA-3 reversibly binds to an inactive PAK conformation, and prevents PAK activation ^20^^,^^21^. Assuming rapid equilibrium of inactive PAK – inhibitor complex, the effect of PAK inhibitor IPA-3 is modelled by considering the concentration of inactive PAK as the following function of [IPA-3],

$$\left[ PAK \right]\left( \left[ IPA\text{-}3 \right] \right)=\frac{\left. PAK \right|_{\left[ IPA\text{-}3 \right]=0}}{\left( 1+\frac{\left[ IPA\text{-}3 \right]}{K_{I}^{PAK}} \right)}$$

(10).

Both, ATP competitive ROCK inhibitor Y‑27632 and ATP bind to an active conformation of the ROCK kinase ^22^^,^^23^. Thus when Y‑27632 is present, the decrease in the ROCK kinase activity can be described by the following multiplier, $\beta<1$,

$$\beta=\left( 1+\frac{\left[ ATP \right]}{K_{d}^{ATP}} \right)/\left( 1+\frac{\left[ ATP \right]}{K_{d}^{ATP}}+\frac{\left[ Y\text{-}27632 \right]}{K_{I}^{ROCK}} \right)$$

(11).

***Dimensionless equations***

To reduce the number of parameters, we express the PDE system, Eq. 9, in a dimensionless form, Eq. 10 ^24^. To simplify the interpretation of numerical results, we left the time as the only dimensional variable (measured in seconds) that directly corresponds to the time, measured in experiments.

| $\frac{\partial rho}{\partial t}=v_{GEF}^{Rho}\frac{1+{\gamma_{DIA}^{Rho}dia}/{k_{DIA}^{Rho}}}{1+{dia}/{k_{DIA}^{Rho}}}\frac{1+{\gamma_{PAK}^{Rho}pak}/{k_{PAK}^{Rho}}}{1+{pak}/{k_{PAK}^{Rho}}}\frac{{rhod}/{k_{GEF}^{Rho}}}{1+{rhod}/{k_{GEF}^{Rho}}}-v_{GAP}^{Rho}\frac{{rho}/{k_{GAP}^{Rho}}}{1+{rho}/{k_{GAP}^{Rho}}}$  $-\nabla\left( -d_{Rho}\nabla rho \right)$  $\frac{\partial rhod}{\partial t}=-v_{GEF}^{Rho}\frac{1+{\gamma_{DIA}^{Rho}dia}/{k_{DIA}^{Rho}}}{1+{dia}/{k_{DIA}^{Rho}}}\frac{1+{\gamma_{PAK}^{Rho}pak}/{k_{PAK}^{Rho}}}{1+{pak}/{k_{PAK}^{Rho}}}\frac{{rhod}/{k_{GEF}^{Rho}}}{1+{rhod}/{k_{GEF}^{Rho}}}+v_{GAP}^{Rho}\frac{{rho}/{k_{GAP}^{Rho}}}{1+{rho}/{k_{GAP}^{Rho}}}$  $-\nabla\left( -d_{RhoD}\nabla rhod \right)$  $\frac{\partial dia}{\partial t}=v_{a}^{DIA}\frac{1+{\gamma_{Rho}^{DIA}rho}/{k_{Rho}^{DIA}}}{1+{rho}/{k_{Rho}^{DIA}}}\frac{\left( d\left( \vec{X} \right)-dia \right)/{k_{a}^{DIA}}}{1+\left( d\left( \vec{X} \right)-dia \right)/{k_{a}^{DIA}}}-v_{i}^{DIA}\frac{{dia}/{k_{i}^{DIA}}}{1+{dia}/{k_{i}^{DIA}}}$  $\frac{\partial rock}{\partial t}=v_{a}^{ROCK}\frac{1+{\gamma_{Rho}^{ROCK}rho}/{k_{Rho}^{ROCK}}}{1+{rho}/{k_{Rho}^{ROCK}}}\frac{\left( r\left( \vec{X} \right)-rock \right)/{k_{a}^{ROCK}}}{1+\left( r\left( \vec{X} \right)-rock \right)/{k_{a}^{ROCK}}}-v_{i}^{ROCK}\frac{{rock}/{k_{i}^{ROCK}}}{1+{rock}/{k_{i}^{ROCK}}}$  $\frac{\partial rac}{\partial t}=v_{GEF}^{Rac}\frac{1+{\gamma_{DIA}^{Rac}dia}/{k_{DIA}^{Rac}}}{1+{dia}/{k_{DIA}^{Rac}}}\frac{1+{\gamma_{PAK}^{Rac}pak}/{k_{PAK}^{Rac}}}{1+{pak}/{k_{PAK}^{Rac}}}\frac{{racd}/{k_{GEF}^{Rac}}}{1+{racd}/{k_{GEF}^{Rac}}}$  $-v_{GAP}^{Rac}\frac{1+{\gamma_{ROCK}^{Rac}\beta rock}/{k_{ROCK}^{Rac}}}{1+\beta{rock}/{k_{ROCK}^{Rac}}}\frac{{rac}/{k_{GAP}^{Rac}}}{1+{rac}/{k_{GAP}^{Rac}}}-\nabla\left( -d_{Rac}\nabla rac \right)$  $\frac{\partial racd}{\partial t}=-v_{GEF}^{Rac}\frac{1+{\gamma_{DIA}^{Rac}dia}/{k_{DIA}^{Rac}}}{1+{dia}/{k_{DIA}^{Rac}}}\frac{1+{\gamma_{PAK}^{Rac}pak}/{k_{PAK}^{Rac}}}{1+{pak}/{k_{PAK}^{Rac}}}\frac{{racd}/{k_{GEF}^{Rac}}}{1+{racd}/{k_{GEF}^{Rac}}}$  $+v_{GAP}^{Rac}\frac{1+{\gamma_{ROCK}^{Rac}\beta rock}/{k_{ROCK}^{Rac}}}{1+\beta{rock}/{k_{ROCK}^{Rac}}}\frac{{rac}/{k_{GAP}^{Rac}}}{1+{rac}/{k_{GAP}^{Rac}}}-\nabla\left( -d_{RacD}\nabla racd \right)$  $\frac{\partial pak}{\partial t}=v_{a}^{PAK}\frac{1+{\gamma_{Rac}^{PAK}rac}/{k_{Rac}^{PAK}}}{1+{rac}/{k_{Rac}^{PAK}}}\frac{\left( p-pak \right)/\left( k_{a}^{PAK}\left( 1+I_{PAK} \right) \right)}{1+\left( p-pak \right)/\left( k_{a}^{PAK}\left( 1+I_{PAK} \right) \right)}-v_{i}^{PAK}\frac{{pak}/{k_{i}^{PAK}}}{1+{pak}/{k_{i}^{PAK}}}$  $rho=\frac{\left[ Rho\text{-}T \right]}{Rho^{tot}}$, $rhod=\frac{\left[ Rho\text{-}D \right]}{Rho^{tot}}$, $rac=\frac{\left[ Rac\text{-}T \right]}{{Rac}^{tot}}$, $racd=\frac{\left[ Rac\text{-}D \right]}{{Rac}^{tot}}$, $pak=\frac{\left[ pPAK \right]}{{PAK}^{tot}}$  $dia=\frac{\left[ DIA^{*} \right]}{{DIA}^{tot}}$, $rock=\frac{\left[ {ROCK}^{*} \right]}{{ROCK}^{tot}}$, $p=\frac{{PAK}^{tot}}{{PAK}^{total}}$, $\vec{X}=\vec{x}/L$, $I_{PAK}=\frac{\left[ IPA\text{-}3 \right]}{K_{I}^{PAK}}$  $\beta=\left( 1+\frac{\left[ ATP \right]}{K_{d}^{ATP}} \right)/\left( 1+\frac{\left[ ATP \right]}{K_{d}^{ATP}}+I_{ROCK} \right)$, $I_{ROCK}=\frac{\left[ Y\text{-}27632 \right]}{K_{I}^{ROCK}}$  $d_{Rho}=\frac{D_{RhoT}}{L^{2}}$, $d_{RhoD}=\frac{D_{RhoD}}{L^{2}}$, $d_{Rac}=\frac{D_{RacT}}{L^{2}}$, $d_{RacD}=\frac{D_{RacD}}{L^{2}}$  $d\left( x \right)=\left( d_{h}-d_{l} \right)\cdot X+d_{l}$, $r=\left\{ \begin{aligned} r_{l}, 0\leq X\leq X_{l} \\ r_{h}, X_{l}\leq X\leq1 \end{aligned} \right.$  $v_{Y}^{X}=V_{Y}^{X}/X^{tot}$, $X=Rho, DIA, ROCK, Rac, PAK$, $Y=GEF, GAP, a, i$  $k_{Y}^{X}=K_{Y}^{X}/X^{tot}$, $X=Rho, DIA, ROCK, Rac, PAK$, $Y=Rho, DIA, ROCK, Rac, PAK, GEF, GAP, i, a$ | (12). |
| --- | --- |

The parameters are listed in Table S2.

**Supplemental Figure Legends**

**Figure S1 related to Figure 1. Elucidation of the topology of the RhoA GTPase network: DIA knockdown influence on the GTPase activities.**

(**A, B**) Pulled-down active forms of RhoA (RhoA-GTP) and Rac1 (Rac1-GTP) and the abundances of DIA, RhoA and Rac1 were measured by Western blot in cells transfected with non-targeting siRNA (control) or siRNA against DIA1 (siDIA1). GST beads were used as negative control, and lipopolysaccharide (LPS) treatment known to activate RhoA was used as positive control for the RhoA-GTP pull-down assay. Numbers 1, 2 and 3 indicate biological replicates. GAPDH was used as loading control. Quantified abundances of RhoA and Rac1 and their relative activities (normalized by the abundances) are shown by bar plots, Asterisks (*) indicate p < 0.05 calculated using unpaired t-test. (**C**) Schematic diagram of positive and negative influences in the RhoA-Rac1 network.

**Figure S2 related to Figure 2. Distinct dynamic regimes of the RhoA-Rac1 network for different effector abundances.**

Domains of the distinct RhoA-Rac1 dynamics partition the planes of the abundances of (**A**) ROCK and DIA, (**B**) PAK and DIA, (**C**) PAK and ROCK. (**D**) Steady-state RhoA-GTP and Rac1-GTP dependences on the ROCK abundance are presented along a 1-D section of the PAK, ROCK plane ($\left[ PAK \right]/\left[ PAK^{tot} \right]=p=0.3$, Eq. 12) shown by white dashed line in (**C**). The amoeboid shape characterized by high RhoA activity emerges in mono-stable region 6 corresponding to the high RhoA-GTP and low Rac1-GTP when ROCK is non-inhibited. (**E**) Wiring diagram of the RhoA-Rac1 signaling network where the dimensionless multipliers, $\alpha_{Y}^{X}$, specify the regulatory influence of protein *Y* on protein *X* (see Star*Methods for details).

**Figure S3 related to Figure 2. Nullclines and vector fields describing the nine dynamic regimes of RhoA-GTP and Rac1-GTP shown in Fig. 2A.**

(**A-I**) Nullclines and vector fields are calculated for a 2-D system given by Eq. 12 for regimes 0 – 8, as indicated. The RhoA-GTP and Rac1-GTP nullclines are shown by red and blue curves, respectively. Projections of limit cycles of a 5-D system in Eq. 6 into a 2-D space of the RhoA and Rac1 activities are shown by green curves. Circles show stable steady states; triangles represent unstable steady states.

**Figure S4 related to Figure 3.**

(**A, B**) Model-predicted snapshots of the RhoA and Rac1 activities for the final phase of the cell movement cycle, when the RhoA-Rac1 wave has reached the cell rear. (**C**) A different biological replicate for the RhoA activity measured in space and time using the RhoA FRET probe during a protrusion-retraction phase (compare to Fig. 3H). (**D**) Spatiotemporal pattern of the RhoA activity during the rear contraction. (**E**) A different biological replicate of a super-resolution fluorescent microscopy image of Rac1 activity (red), combined with staining for F-actin (phalloidin, white) and the nucleus (DAPI, blue) in fixed cells. The image shows the Rac1 activity wave propagation into the cell body. (**F**) Computational mesh used in calculations for a given cell shape.

**Figure S5 related to Figure 4.**

(**A, B**) Model-predicted dependencies of the RhoA and Rac1 activities on the concentration of PAK inhibitor ($I^{PAK}=\left[ IPA\text{-}3 \right]/K_{I}^{PAK}$) for gradually increasing (blue) and decreasing (red) PAK inhibitor concentrations. The network evolution occurs through two different routes (blue and red curves in Figs. S5A and S5B). It is calculated by averaging the GTPase activities over time and the cell volume based on Western blot measurements in our previous work ^6^.

**Figure S6 related to Figure 5.**

(**A**) A different biological replicate of fluorescent microscopy image of Rac1 activity (red), and F-actin (phalloidin, white) and nuclear (DAPI, blue) staining in fixed MDA-MB-213 cells treated with 2.5µM Y-27632 ROCK inhibitor for 15 min. (**B**) Model-predicted numbers of RhoA activity bursts during 10 minutes at the leading edge and the rear for control cells and cells where ROCK was inhibited by 2.5 μM of Y‑27632. (**C**) The number of experimentally observed RhoA activity bursts during 10 min measured using the RhoA FRET probe at the leading edge and the rear for control cells and cells treated with the ROCK inhibitor Y‑27632. Error bars represent 1^st^ and 3^rd^ quartiles, asterisks *** indicate p < 0.001 calculated using unpaired t-test. (**D**) Quantification of the total distance travelled by the control cells and cells treated with a ROCK inhibitor. Results were similar for two different ROCK inhibitors, 10 μM Y‑27632 and 1 μM GSK269962A. (**E**) Model predicted lower threshold of DIA abundance for induction of RhoA-Rac1 oscillations (region 1 in Fig. 2A) in control and DIA knockdown conditions. DIA knockdown conditions were modeled by changing of RhoA and Rac1 abundance according to experimental data (Figs. S1A,B).

**Supplemental Tables**

| **Table S1.** **Quantitative mass spectrometry data** **^6^** **used to populate protein abundances in our mathematical model.** | | | |
| --- | --- | --- | --- |
| **Gene name** | **Protein name** | **Copy number intensity** | **Concentration (nM)*** |
| PAK2 | Serine/threonine-protein kinase PAK 2 | 661552 | 27.00 |
| RAC2 | Ras-related C3 botulinum toxin substrate 2 | 277301 | 11.09 |
| RAC1 | Ras-related C3 botulinum toxin substrate 1; Ras-related C3 botulinum toxin substrate 3 | 3984262 | 160.00 |
| RHOT1 | Mitochondrial Rho GTPase 1;Mitochondrial Rho GTPase | 7391 | 0.30 |
| RHOC | Rho-related GTP-binding protein RhoC | 530247 | 21.21 |
| RHOA | Transforming protein RhoA;Rho-related GTP-binding protein RhoC | 4013548 | 161.00 |
| RHOG | Rho-related GTP-binding protein RhoG | 490041 | 19.60 |
| RHOT2 | Mitochondrial Rho GTPase 2 | 7850 | 0.31 |
| RHOF | Rho-related GTP-binding protein RhoF | 25597 | 1.02 |
| ROCK1 | Rho-associated protein kinase 1 | 13858 | 0.55 |
| ROCK2 | Rho-associated protein kinase 2 | 62134 | 2.49 |
| DIAPH1 | Protein diaphanous homolog 1;Diaphanous homolog 1 (Drosophila), isoform CRA_a | 189210 | 7.57 |
| DIAPH2 | Protein diaphanous homolog 2;Diaphanous homolog 2 (Drosophila), isoform CRA_c;Diaphanous homolog 2 (Drosophila), isoform CRA_a | 8292 | 0.33 |
| DIAPH3 | Protein diaphanous homolog 3 | 19067 | 0.76 |

^*^ Concentrations were calculated assuming an average cell volume $4\cdot{10}^{-14}$ L.

| **Table S2.** **Parameter values (Eq. 12).** | | | | |
| --- | --- | --- | --- | --- |
| **Parameter** | **Value** |  | **Parameter** | **Value** |
| $\gamma_{Rho}^{DIA}$ | 5.0526 |  | $k_{a}^{DIA}$ | 2 |
| $\gamma_{Rho}^{ROCK}$ | 22.7795 |  | $k_{a}^{ROCK}$ | 2 |
| $v_{a}^{DIA}$ | 19.2456 1/s |  | $v_{a}^{PAK}$ | 0.0186 1/s |
| $v_{i}^{DIA}$ | 10.5262 1/s |  | $k_{a}^{PAK}$ | 0.288 |
| $k_{i}^{DIA}$ | 0.0158 |  | $v_{i}^{PAK}$ | 0.089 1/s |
| $v_{a}^{ROCK}$ | 0.2470 1/s |  | $k_{i}^{PAK}$ | 0.16 |
| $v_{i}^{ROCK}$ | 9.5818 1/s |  | $\gamma_{DIA}^{Rho}$ | 100 |
| $k_{i}^{ROCK}$ | 0.0395 |  | $k_{DIA}^{Rho}$ | 3 |
| $v_{GEF}^{Rho}$ | 0.4902 1/s |  | $\gamma_{DIA}^{Rac}$ | 7.8 |
| $k_{GEF}^{Rho}$ | 0.3591 |  | $k_{DIA}^{Rac}$ | 0.055 |
| $v_{GAP}^{Rho}$ | 0.7707 1/s |  | $\gamma_{ROCK}^{Rac}$ | 10 |
| $k_{GAP}^{Rho}$ | 0.0218 |  | $k_{ROCK}^{Rac}$ | 0.05 |
| $v_{GEF}^{Rac}$ | 0.1118 1/s |  | $k_{Rho}^{DIA}$ | 0.04 |
| $k_{GEF}^{Rac}$ | 0.0275 |  | $k_{Rho}^{ROCK}$ | 1.3 |
| $v_{GAP}^{Rac}$ | 0.5495 1/s |  | $\gamma_{PAK}^{Rho}$ | 0.025 |
| $k_{GAP}^{Rac}$ | 0.0109 |  | $k_{PAK}^{Rho}$ | 0.012 |
| $\gamma_{Rac}^{PAK}$ | 6.7 |  | $k_{Rac}^{PAK}$ | 0.65 |
| $\gamma_{PAK}^{Rac}$ | 1 |  | $k_{PAK}^{Rac}$ | 0.1 |
| $d_{Rac}$ | 0.0005 1/s |  | $d_{Rho}$ | 0.0005 1/s |
| $d_{RacD}$ | 0.0005 1/s |  | $d_{RhoD}$ | 0.0005 1/s |
| $d_{h}$ | 1.1 |  | $d_{l}$ | 0.8 |
| $r_{h}$ | 1.85 |  | $r_{l}$ | 0.5 |
| $X_{l}$ | 0.8 |  | $p$ | 1 |
| $\left[ ATP \right]/K_{d}^{ATP}$ | 112 |  | $K_{I}^{ROCK}$ | 220 nM |

| **Table S3. Key experimental and computational resources.** | | |
| --- | --- | --- |
| **REAGENT or RESOURCE** | **SOURCE** | **IDENTIFIER** |
| **Antibodies** | | |
| Anti-Rac1 clone 23A8 | Millipore | cat.05-389 |
| Anti-RhoA (26C4) | Santa-Cruz Biotechnology | cat.sc-418 |
| anti-GAPDH (D16H11) XP® | CST | cat.5174 |
| anti-DIA1 | Thermo | cat.PA5-21409 |
| anti-ROCK1 | Thermo | cat.PA5-22262 |
| anti-Rac-GTP | New East Bio | cat.26903 |
| Anti-mouse F(ab')2 Fragment Alexa Fluor ® 647 | Thermo | cat.. A-21237 |
| Anti-rabbit F(ab')2 Fragment Alexa Fluor ® 594 | Thermo | cat. A-11072 |
| Anti-rabbit Alexa Fluor-488® | Thermo | cat. A-21206 |
| Anti-rabbit Alexa Fluor-594® | Thermo | cat. A-11012 |
| Anti-rabbit IgG, HRP-linked | CST | cat.7074 |
| Anti-mouse IgG, HRP-linked | CST | cat.7076 |
| **Bacterial and Virus Strains** | | |
| IncuCyte® NucLight Red Lentivirus Reagent | Essen | Cat. 4625 |
| **Chemicals, Peptides, and Recombinant Proteins** |  |  |
| Y-27632 | Sigma Aldrich | Cat.Y0503 |
| GSK 269962 | Selleckchem | Cat.S7687 |
| GST-Beads | Sigma Aldrich | G.4510 |
| 4,6-Diamidino-2-phenylindole dihydrochloride (DAPI) | Sigma Aldrich | Cat.10236276001 |
| Rhodamine Phalloidin | Thermo | Cat. R415 |
| Phalloidin- Alexa Fluor-488 | Thermo | Cat. A12379 |
| VECTASHIELD antifade mounting media | Vector labs | Cat. H-1000 |
| Dulbecco’s Modified Eagle Medium (DMEM) | Sigma Aldrich | Cat.D6429 |
| FluoroBrite DMEM Media | Thermo | Cat. A1896701 |
| Fetl Bovine Serum (FBS) | Gibco | Cat.10270 |
| Collagen (rat tail) | Sigm Aldrich | Cat.11179179001 |
| Puromycin | Sigma Aldrich | Cat. P8833 |
| Polibrene | Millipore | Cat.TR-1003-G |
| Lipofectamine RNAiMax | Thermo | Cat.13778 |
| **Experimental Models: Cell Lines** |  |  |
| MDA-MB-231 | ATCC | Cat.HTB-26™ |
| **Oligonucleotides** |  |  |
| DIAPH1 siRNA SMART Pool | Dharmacon | cat. L-010347-00-0010 |
| **Recombinant DNA** |  |  |
| GST-rhotekin-RBD | Dr. Mike Olson gift (Beatson Institute, Glasgow, UK) |  |
| GST-PAK-CRIB | Dr. Piero Crespo gift (IBBTEC,University of Cantanbria, Spain) |  |
| mTFP-YFP RhoA activity probe | Prof. Olivier Pertz Gift (Institute of Cell Biology, Bern, Switzerland) |  |
| psPAX-2 | Prof. Olivier Pertz Gift |  |
| VsVg | Prof. Olivier Pertz Gift |  |
| **Software and Algorithms** |  |  |
| Fiji | ^25^ | https://imagej.net/Fiji |
| OpenFOAM | ^26^ | https://www.openfoam.com/ |
| ParaView | ^27^ | https://www.paraview.org/ |
| Salome | ^28^ | https://www.salome-platform.org/ |
| Python |  | https://www.python.org/ |
| SciPy | ^29^ | https://www.scipy.org/ |
| MatplotLib | ^15^ | https://matplotlib.org/ |
| OpenCV | ^30^ | https://opencv.org/ |
| DYVIPAC | ^13^ | https://bitbucket.org/andreadega/dyvipac-python/src/master/ |
| BioNetGen | ^11^^,^^12^ | https://www.csb.pitt.edu/Faculty/Faeder/?page_id=409 |

**Supplemental Videos**

**Video S1.** Model-predicted spatiotemporal activity patterns of RhoA and Rac1.

**Video S2.** Live-cell imaging of cel movement cycles. Red color represents staining of the nuclei.

**Video S3.** Model-predicted spatiotemporal activity patterns of RhoA and Rac1 when ROCK is inhibited by 2.5 μM of Y‑27632.

**Video S4.** Live-cell imaging of cellular movement cycles in cells treated with 10 μM Y‑27632 ROCK inhibitor. Red color represents staining of the nuclei.
