## Supplementary figures and images for "Periodic propagating waves coordinate RhoGTPase network dynamics at the leading and trailing edges during cell migration"

### Figure S1

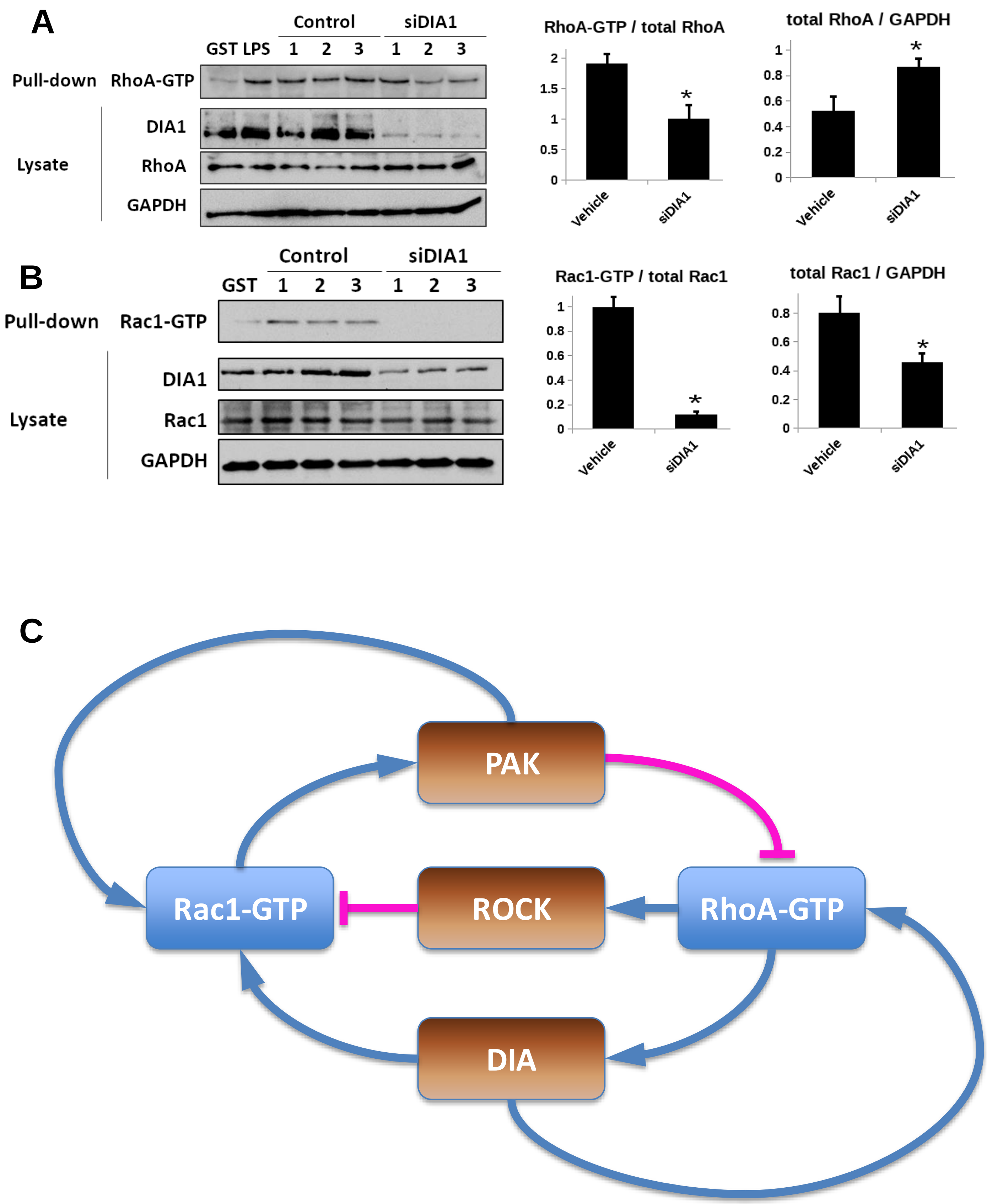

### Figure S2

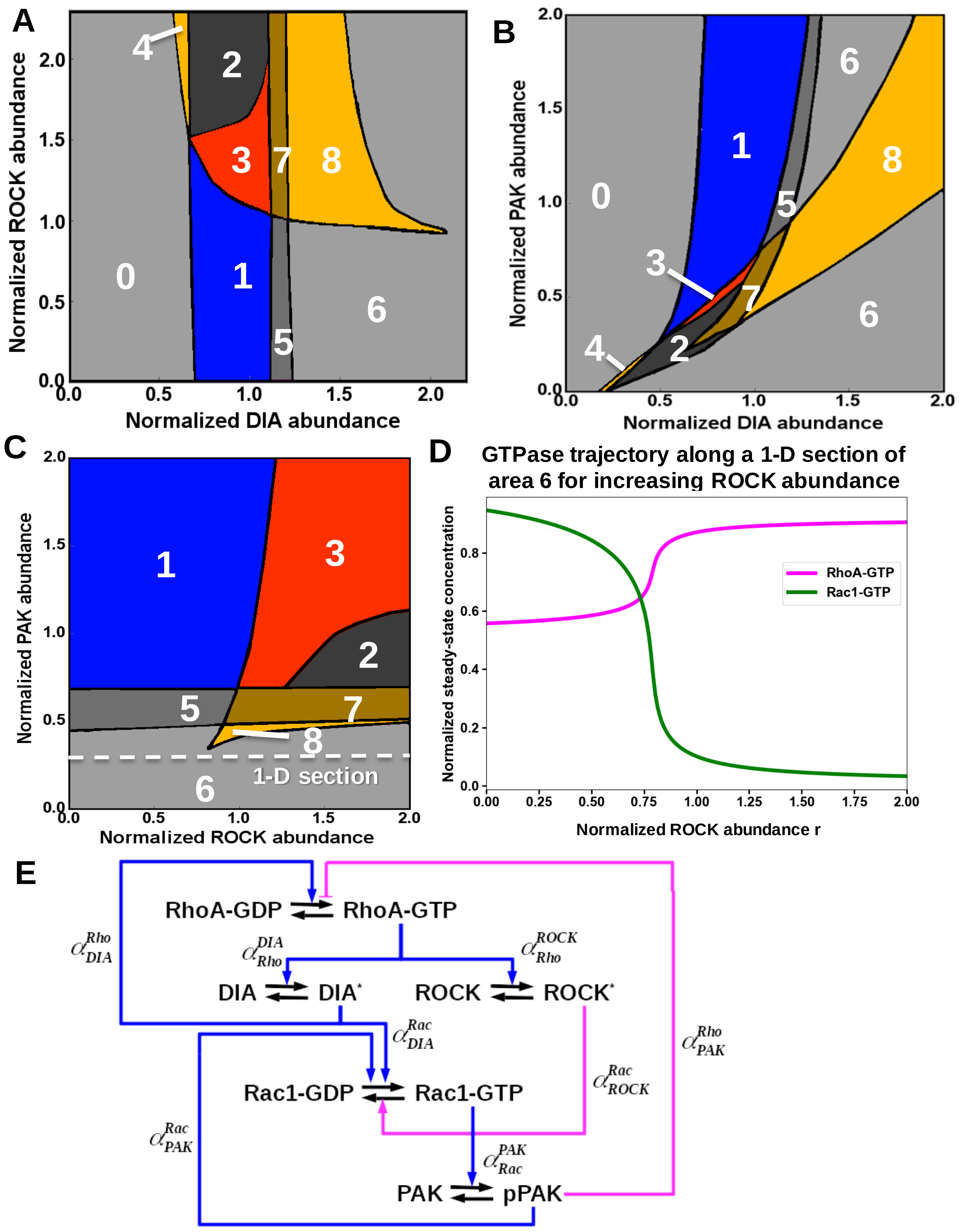

### Figure S3

#### Figure S3

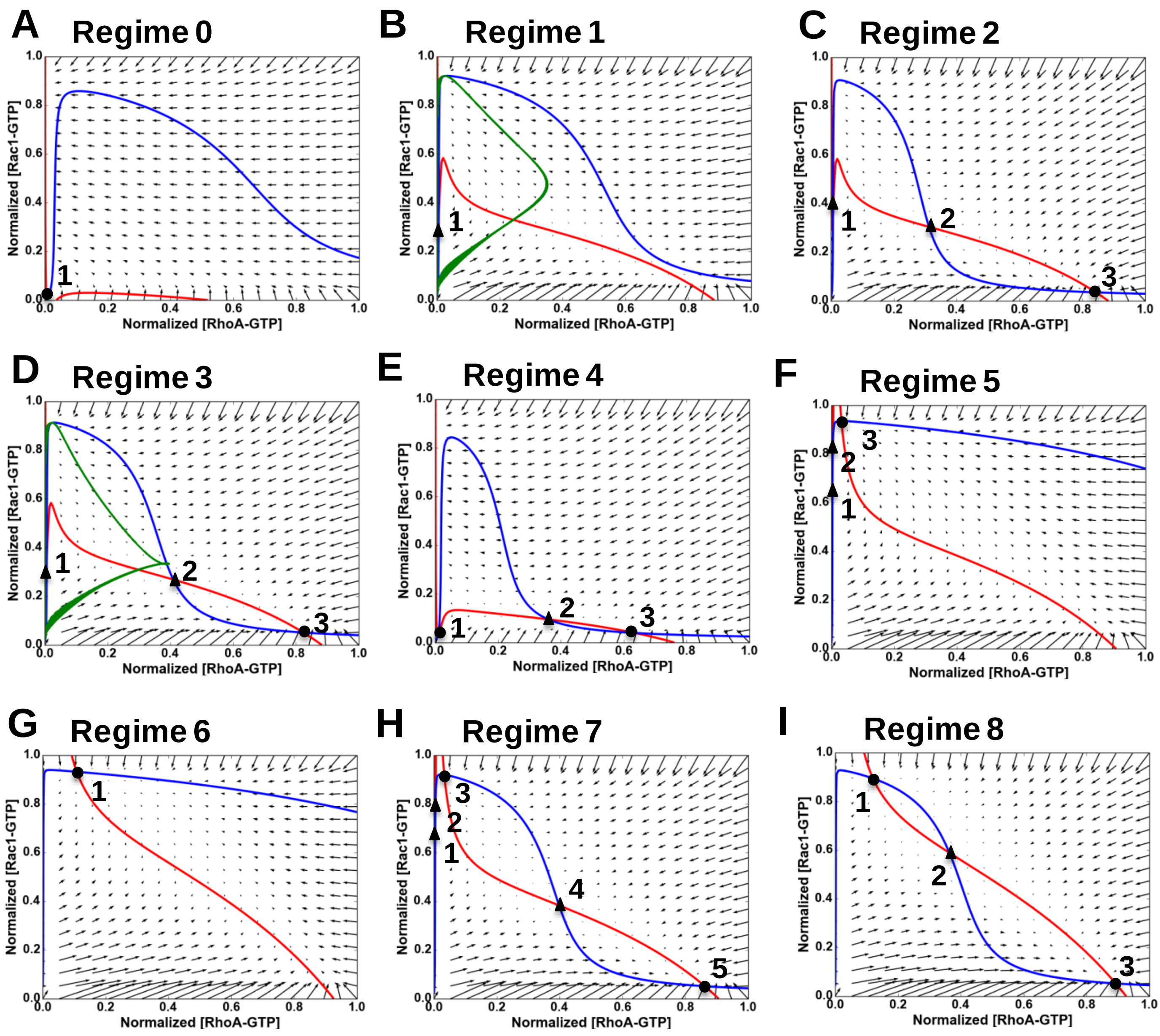


d [Rho-T] / dt = 0

d [Rac-T] / dt = 0

Limit cycle trajectory

### Figure S4

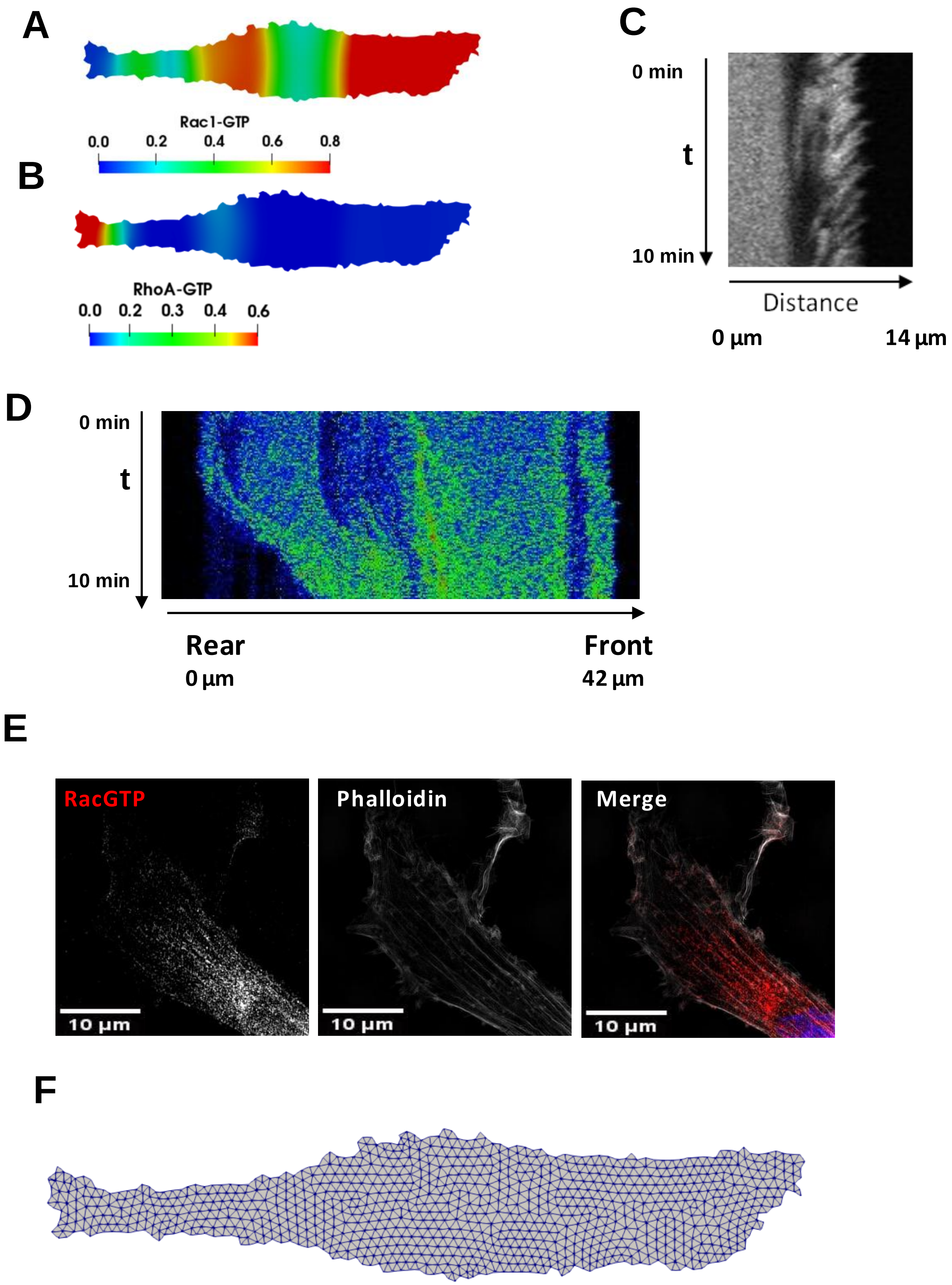

### Figure S5

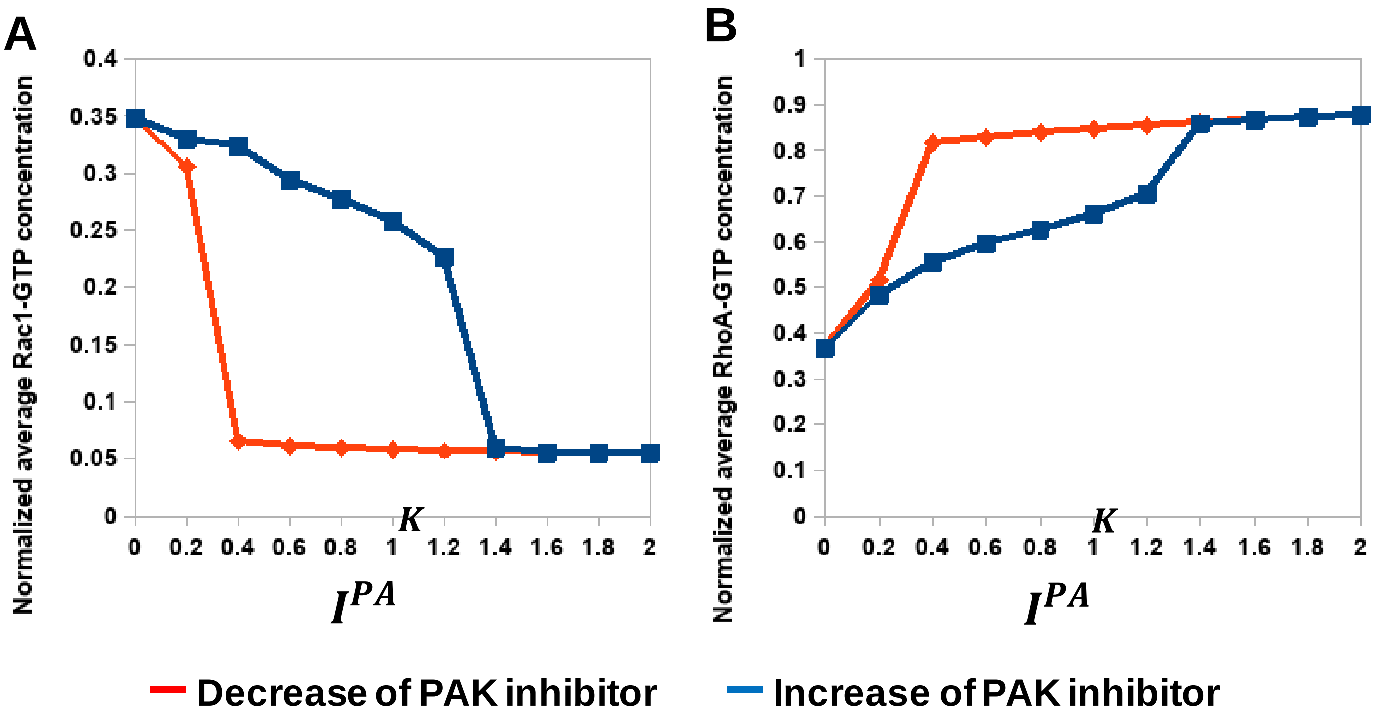

### Figure S6

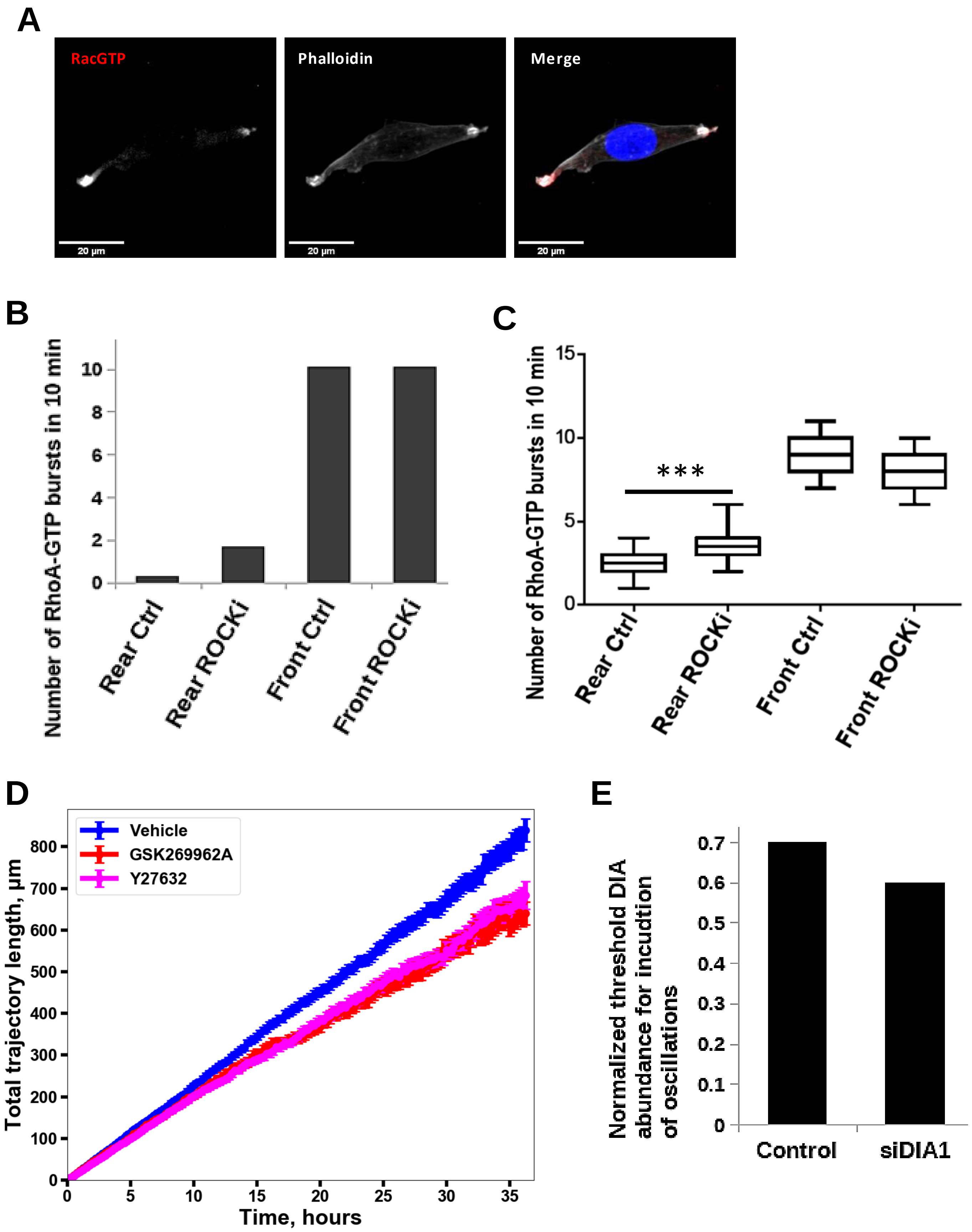
